# The schizophrenia associated protein DISC1 is a multivalent tetrameric hub of conserved ancient fold

**DOI:** 10.64898/2026.01.28.701986

**Authors:** Jin Chuan Zhou, Jiri Kratochvil, Fei Ye, Kamel el Omari, Philipp Kukura, Mingjie Zhang, Elena Seiradake

## Abstract

DISC1 is a pleiotropic protein with essential roles in neuronal proliferation and migration, intracellular signalling and cargo transport. It associates with a diverse array of partner molecules in these contexts. Mutations at the *DISC1* locus are strongly associated with a spectrum of mental illnesses such as schizophrenia and depression. Despite its clinical relevance, the molecular architecture and function of DISC1 have remained largely elusive. We present a cryo-EM structure of the entire conserved core region of DISC1. The structure reveals an intricate homotetrameric assembly that harbours conserved bacteria-derived UVR domains. Four of these domains, one from each monomer, mediate extensive contacts forming two asymmetric dimer units. The dimers in turn interface with each other at two distinct coiled coil domains to achieve a two-fold symmetric tetramer. Mutational analysis shows that this tetrameric architecture enables DISC1 to simultaneously bind multiple copies of NDE1 client protein. Importantly, tetramerization and partner binding are structurally independent functions of DISC1. Altogether, our study provides a compelling molecular model of an ancient bacteria protein fold participating in the assembly of a multivalent mammalian scaffold hub that can coordinate multiple partner molecules.

## Introduction

The Disrupted in Schizophrenia 1 (*DISC1*) is recognised as a prominent susceptibility gene for major mental illnesses such as schizophrenia, bipolar disorder and major depression (1). The *DISC1* locus was initially identified as the target of a balanced translocation event, t(1;11)(q42.1,q14.3), within members of a Scottish family affected by schizoaffective disorders (2–4). Since its discovery, evidence of partial deletion, copy number variants (CNVs) as well as single nucleotide polymorphisms (SNPs) affecting the *DISC1* locus have been reported in psychiatric patients (5–8). Consequently, much effort has undergone to understand the biological context of DISC1 function and to pinpoint the molecular events that underlie the observed clinical phenotypes. The consensus view emerging from hundreds of studies to date portrait DISC1 as a protein of pleiotropic functions, for example, fulfilling important roles in neuron proliferation, neurite outgrowth, centrosome maintenance, intracellular signalling and cargo transport (9–13). These studies have demonstrated that DISC1 is part of an increasingly understood and complex interactome (14,15). Importantly, a number of DISC1 partner proteins such as NDEL1 and GSK3β have also emerged as genetic risk factors associated with psychiatric disorders (16,17), suggesting that their concerted actions play a role in these conditions. The data point to a converging network of interacting players where DISC1 acts as a coordinating hub (14).

One of the best characterised binding partners of DISC1 is NDE1/NDEL1, a regulatory subunit belonging to the dynein motor complex (18). NDE1 is known to form elongated oligomers of simple topology via its coiled-coil domains (CC) (19,20). Previous solution NMR studies have established the presence of a CC motif at the very C-terminal end of DISC1 (DISC1-C, residues 767-829) that is targeted with nanomolar affinity by NDE1 (9) (Fig. 1a). This binding module is tethered to the bulk of DISC1 protein via a low-complexity linker.

**Fig 1.**
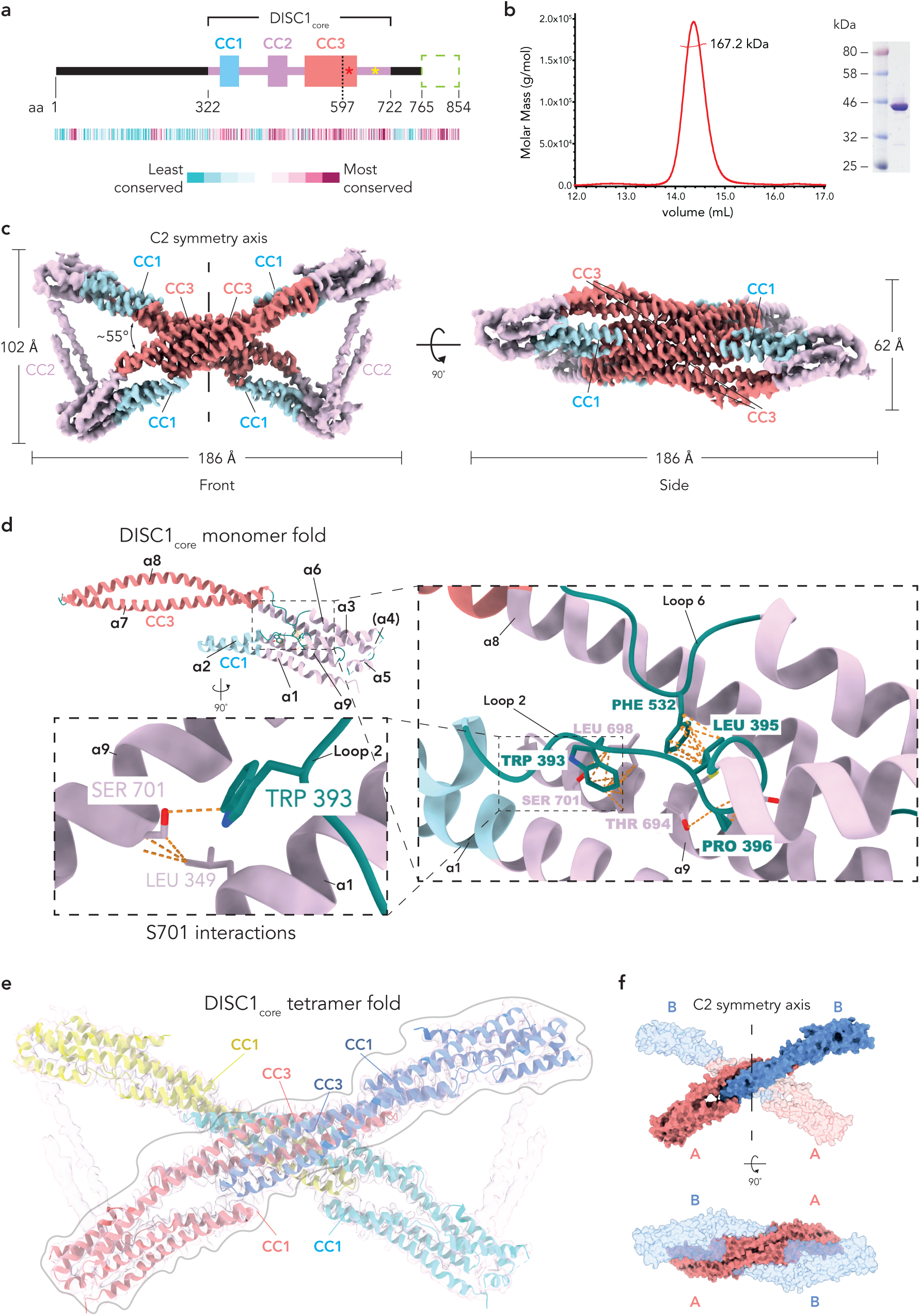
DISC1_core_ forms a homotetramer. **a**: A simplified, overall view of human DISC1 domain architecture. DISC1-N (aa. 1-321) is unstructured while the downstream region denoted as DISC1_core_ (aa. 322-772) is predicted to harbour three coiled coil motifs (colour-coded boxes labelled respectively as CC1, CC2 and CC3). Lying C-terminally to DISC1_core_ is a further coiled coil domain (DISC1-C, aa. 765-854, green dashed box), which is a well-characterised binding site for partner proteins NDE1 and ATF4. The dashed line overlapping CC3 refers to the point of truncation caused by the familial t(1;11) translocation mutation. Sites for the common risk variants L607F and S704C are shown by the red and yellow asterisks respectively. The heatmap below illustrates the amino acid conservation across each protein region, where blue residues are poorly conserved while purple residues show a high degree of evolutionary conservation. **b**: On the left, MALS elution profile showing that DISC1_core_ behaves as a homotetramer in solution. Expected molecular weight for a monomer of DISC1_core_ is ∼ 45 kDa. On the right, Coomassie-stained SDS-PAGE of the DISC1 sample used for the MALS assay. **c**: Cryo-EM map of DISC1_core_ tetramer. Left and right panels show front and side views respectively of the ‘bow tie’ like DISC1 tetrameric fold. The positions of the three assigned CC domains are annotated here. In particular, CC1 and CC3 regions are coloured in light blue and coral red respectively. The overall dimensions of the protein complex, the two-fold symmetry axis as well as the angle of intersection between the two constituent dimers are indicated. **d**: Atomic model of a single DISC1_core_ monomer. The position for each of the 9 helices in DISC1_core_ is annotated, including that of α4 (in bracket), the only helix that is not present in our structure. CC1 and CC3 regions are coloured as in c. Large right inset provides a close-up view of the two extended loops 2 and 6, which are anchored against helix α9 to help maintain an overall compact monomer fold. The conserved hydrophobic residues that are key to loop stabilisation are highlighted here. All loop regions are coloured in teal green. Lower left inset provides a close-up view depicting the structural role played by S701 (equivalent to S704 in human DISC1) which contacts both W393 in loop 2 and L349 in α1. The orientation of the inset view with respect to the reference structure is also indicated. **e**: Atomic model of DISC1_core_ tetramer. The model is shown here fitted into the EM map with each protomer color-coded. The positions for all four copies of CC1 as well as for two of the four CC3 are annotated. A light grey contour highlights one of the two contributing dimers in the structure. **f**: DISC1_core_ tetramer exists as a ‘dimer of dimers’. Front and side view (top and bottom surface representations, respectively) showing how the two-fold symmetry of DISC1_core_ is dictated by the asymmetric nature of the AB dimer, consisting of the two structurally distinct A and B forms of monomer (coloured in coral red and cornflower blue, respectively). The bottom panel also clearly indicates that the two A protomers help to bridge the dimers into a tetramer.

To date, this small DISC1-C domain is the only structurally defined part of the protein while the molecular architecture for the majority of DISC1 remains unknown. *In silico* sequence analysis predicts a string of three CC motifs (CC1 to CC3) upstream of the DISC1-C linker (Fig 1a, Supp Fig. 1a). This middle, or ‘core’ segment coincides with a high degree of evolutionary conservation (encompassing residues 321-725). Previous sequence-based bioinformatic searches uncovered two conserved, homologous repeats within the primary sequence of this region (21). Surprisingly, these repeats present striking similarities to the UVR domain found in bacterial proteins such as UvrB and UvrC in *E.coli*, which have a role in repairing UV-induced DNA damage (22,23). Albeit not strictly limited to the prokaryotic kingdom, the UVR domain is known to adopt a conserved structure consisting of two short helices packed into an antiparallel hairpin fold (24). The amino acid contacts along the two interfacing helices typically follow the pattern of a coiled coil. A common feature shared by several UVR domain proteins is their ability to dimerise via a ‘head to head’ contact involving the central linker within the UVR domain hairpin (24,25). Whether such an arrangement is important in DISC1 remains unexplored.

In stark contrast to the central domain, an extensive segment of DISC1 N-terminal region (DISC1-N, covering residues 1-320) retains poor sequence conservation among vertebrates (26), with the first two exons sharing only about 50% identity between mouse and human (21). The local sequence here is unusually enriched in serine and glycine residues (21,26), thus showing a high propensity towards protein disorder.

To understand the molecular architecture and functional significance of the highly conserved DISC1 core region, we determined the cryo-electron microscopy (cryo-EM) structure of DISC1_core_, revealing an unexpectedly elaborate, homotetrameric fold. Key to this assembly are two conserved UVR domains that define an asymmetric ‘dimer of dimers’ packing of the four DISC1 monomers. We further show that the tetramer architecture enables simultaneous binding of multiple NDE1 client molecules by DISC1. Importantly, DISC1 tetramerisation and binding to NDE1 are structurally independent functions. Altogether, our findings confirm DISC1 function as a scaffolding hub that spatially coordinates its many partners.

## Results

### DISC1 harbours a conserved tetrameric core

#### DISC1 tetramer adopts an intricate helical architecture

Previous efforts to understand the molecular structure of DISC1 have been impeded by the intractable nature of the purified full-length protein *in vitro* (27,28). Here, we identified a highly conserved, uncharacterised central domain of the murine DISC1 homologue (residues 322-722) which is amenable to structural investigations. We designate this region as DISC1_core_ (Fig. 1a). Size-exclusion chromatography (SEC) and SDS-PAGE analysis confirmed the homogenous and monodisperse nature of our purified protein fraction (Supp Fig. 1b). Surprisingly, although the predicted molecular mass of this DISC1 sample is ∼45 kDa, measurements by SEC coupled with multi-angle light scattering (SEC-MALS) indicated the formation of a ∼170 kDa homotetrameric complex in solution (Fig. 1b). Therefore, we next sought to determine the molecular structure of this DISC1 assembly via single particle cryo-EM (Supp Fig. 2a-d, Table S1).

A total of ∼227,000 selected particles contributed to the reconstruction of a 3.9Å EM density map (Supp Fig. 2b). In agreement with *in silico* predictions, our map reveals a complex structure consisting almost exclusively of α helices (Fig. 1c). These are intricately intertwined together, which starkly contrasts with the simpler, extended topology shown or predicted for many coiled coil partners of DISC1, such as NDE1 (19). The overall architecture of DISC1_core_ tetramer adopts a distinctive hollow “bow tie” shape, spanning ∼180Å in length and ∼100Å in height (Fig. 1c). The complex arrangement is largely composed of two prominent helical bundles (each attributable to a dimer of DISC1_core_, as detailed below) that are juxtaposed at an 55° angle across. Evidence to the structural stability invoked by this type of helical packing, the local resolution in this area of the map is estimated to surpass 3.5Å (Supp Fig. 2d). Densities corresponding to closely packed helices of varying lengths are therefore clearly discernible (Fig 1c). Importantly, many instances of distinct side chain densities at these packing interfaces can also be observed (Supp Fig. 3a-d). Our map shows the entire DISC1 complex to be centred around a 60Å thick, four-layered stack of partially overlapping coiled coils. These correspond to four copies of CC3 which are the longest identifiable coiled coils here, each stretching ∼ 80Å across the map. In their close proximity lie CC1 densities from each of the monomers, which are well positioned to contact and enclose the CC3 core from four different directions. Thus, CC1s essentially function as buttresses to support and firmly secure their CC3 partners in position. Providing further stability to the overall assembly are partially resolved densities that we assigned to two copies of CC2 at each lateral side. These are integral to the “bow tie” arrangement by linking the four corners of the structure. Altogether, our cryo-EM data confirms the predicted dominant role played by coiled-coil interactions in the assembly of the DISC1 core domain.

Our map enables confident modelling of all 9 α helices that form part of a monomer of DISC1_core_ (Fig. 1d, Supp Fig. 4a, Table S2), except for α4, which belongs to the weak CC2 densities. As alluded to previously, DISC1_core_ assumes a relatively compact arrangement. While α1, α2 and α7, α8 contribute to CC1 and CC3 formation respectively, the extensions in α1 and α8 closely bundle together with the remaining helices. Here, helix α9 plays an important support role. The compactness of the monomer fold is further aided by two extended loops in between α2, α3 (loop 2) and α6, α7 (loop 6) respectively (Fig. 1d, right inset). In particular, a group of well conserved hydrophobic residues, namely W393, L395, F532 and L698, cluster together to create a stable anchoring of the loops against α9. Of note is S701 which is well positioned to bridge α9 to loop 2 and α1 via contact with W393 and L349 respectively (Fig. 1d, bottom inset). Previous studies have shown that a SNP at the S701 equivalent in human DISC1, namely S704C (Fig. 1a), is one of the most common risk variant for schizophrenia found in the population (29).

#### DISC1 tetramer has only two-fold symmetry

Given the homotetrameric nature of DISC1_core_, it is surprising that our EM map reveals an overall two-fold rather than four-fold symmetry, with the rotational symmetry axis falling across the central coiled coil stack (Fig. 1c). To understand the underlying principle for the observed symmetry, we fitted four copies of DISC1 monomer into our map (Fig. 1e). Intriguingly, during the fitting process, we found significant structural differences in two copies of monomers (protomers A) with respect to the other two (protomers B). In each instance, there is a notable shift in tilt of ∼11° in CC3 positioning which is also accompanied by a slight torsion of CC1, together accounting for a measured root-mean-square deviation (RMSD) of 7.5Å between the two A and B forms of protomer (Supp Fig. 4b). Consequently, the DISC1 tetramer complex is best described as a ‘dimer of dimers’, whereby each A and B protomers combine into an asymmetric dimer unit (Fig. 1f, top). The two copies of protomer A then interface with each other, bringing the two dimers together (Fig. 1f, bottom). Thus, DISC1_core_ two-fold symmetry is dictated by the asymmetric nature of its dimer unit, which in turn is influenced by the differential geometries adopted by CC1 and CC3 within A and B protomers.

### CC1 G CC3 are integral elements of the DISC1 fold

#### A conserved role for UVR domain in the asymmetric DISC1 dimer

Our atomic model shows that the orientations of CC1 and CC3 relative to each other, with CC3 protruding over CC1, make them perfectly posed to form a compact dimer. Interestingly, we mapped the two predicted UVR motifs to CC1 (CC1_UVR_, residues 351-389) and part of CC3 (CC3_UVR_, residues 568-625) respectively (Fig. 2a). In accordance with their suggested functional relevance, these domains are located right at the heart of the interface between each A and B protomer. In particular, each CC1_UVR_ interlocks with the CC3_UVR_ of the opposite monomer by exerting the same type of head to head hairpin loop interaction as described for the UVR class of bacterial proteins (24). The conserved nature of the contacts here is remarkable: both in the bacterial UVR archetypes (Supp Fig. 4c) and in mammalian DISC1, a local hydrophobic pocket is created via clustering of residues from opposing loops (DISC1 V366, Y371 and A374 in both CC3_UVR_ packing against L593, F598 in the two CC1_UVR_, Fig. 2a left and right insets). This is also reminiscent of the hydrophobic core observed at the interface of other UVR domain proteins, such as the Clp chaperones in plants chloroplast systems (25). Further evolutionary analysis shows that this dimerization mechanism likely extends to other DISC1 homologues that are also known to harbour two or more copies of UVR domains (Fig. 2b) (21). Indeed, structural comparison of AlphaFold predicted homodimers for several DISC1 homologues across main eukaryotic phyla such as Opisthokonta, Plantae and Excavates shows retention of the same UVR dimer architecture with key interface residues such as Y371 and F598 being absolutely conserved (Fig. 2b right upper panel inset). Testament to the critical role performed by this hydrophobic core, mutations targeting only four of the residues in the region (including Y371 and F598, DISC1 ΔN_mut C_, Table S3) are sufficient to completely abrogate DISC1 tetramer formation (Supp Fig. 5, middle plot, Table S4).

**Fig 2.**
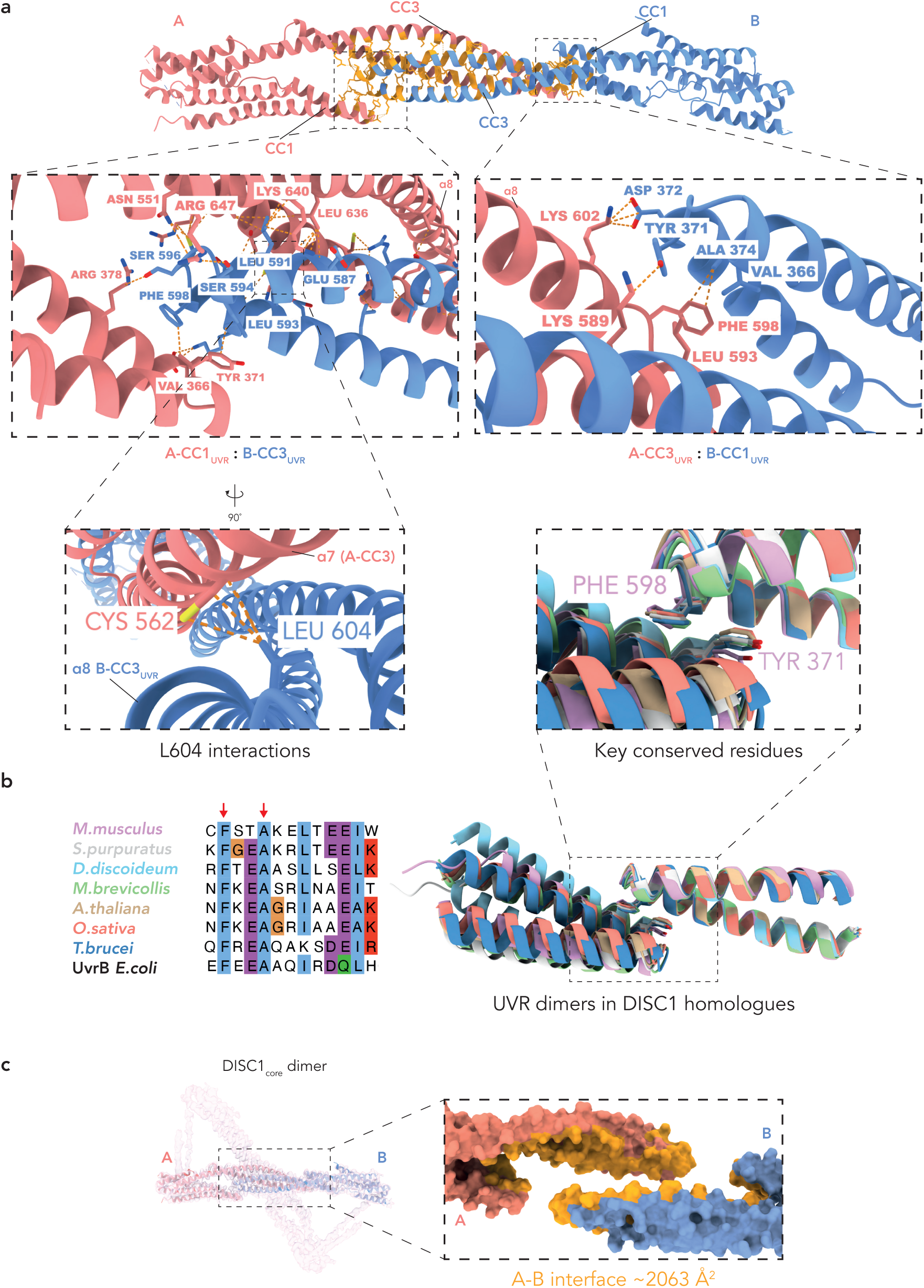
UVR domains in CC1 and CC3 are key to DISC1_core_ dimerization. **a**: Overview of all contacting residues (in orange) at the A-B (‘monomer-monomer’) interface, clearly depicting the dominant role of CC1 and CC3 in mediating DISC1 dimerization. The two copies of CC1 and CC3 are indicated. Each monomer is coloured as in Fig 1f. Below are close-up views illustrating the structural function of UVR domains in both CC1 and CC3. In particular, left and right insets highlight the bias in contacting residues for A-CC1_UVR_: B-CC3_UVR_ binding (left) compared to A-CC3_UVR_: B-CC1_UVR_ binding (right). Lower inset, close-up view of the A-CC1_UVR_: B-CC3_UVR_ binding site focusing on the contribution of L604 (equivalent to L607 in human DISC1) to the A-B interface. **b**: Left, multiple sequence alignment encompassing one of the two UVR domains for several known DISC1 homologues: *M.musculus* (Q811T9), *S.purpuratus* (UPI0000E48D70), *D.discoideum* (Q550K0), *M.brevicollis* (A9UYG7), *A.thaliana* (Q8L4Q6), *O.sativa* (Q84Z60) and *T.brucei* (Q57ZU7). Last sequence corresponds to the UVR domain from the *E.coli* UvrB (P0A8F8). Two extremely conserved interface residues F598 and A601 are highlighted here with red arrows. Sequence alignment was generated with MAFFT (55). Right, structural superposition of DISC1 homologues colour-coded as per legend in the left panel. Upper inset, close-up view of the conserved UVR dimer interface. For clarity, only two sets of the conserved hydrophobic residues, equivalent to F598 and Y371 in murine DISC1, are depicted here. **c**: One asymmetric AB dimer is fitted into the EM map. Left inset portraits an exploded view of the surface representation for the A-B interface, highlighting the buried surface area, shown in orange here.

Highlighting the asymmetric nature of DISC1_core_ A and B protomers, the two pairs of CC1_UVR_: CC3_UVR_ do not contribute equally to dimer formation. Because of CC3’s coiling geometry, the two CC3_UVR_ motifs do not approach their respective target monomer with the same orientation, but rather with a roughly 90° angular difference (Fig. 2a left vs right inset). As a consequence, in both dimers, it is always the CC3_UVR_ head from protomer B that is favourably positioned for insertion into the cavity created between CC1_UVR_ and the end region of CC3 of protomer A (Fig. 2a left inset). Here, the hydrophobic packing of the helices is reinforced by additional hydrogen bonds and salt bridges (occurring between D372, N551, C562, K566, K640, R647 and their respective targets K602, G595, E603, E607, E587, S594) effectively locking the CC3_UVR_ of protomer B into the cleft within protomer A. Part of B-CC3_UVR_ and also contributing to this interface is L604, which contacts C562 on α7 of the opposing A-CC3 (Fig. 2a, lower inset). Notably, a SNP at position L607 in human DISC1 (Fig. 1a) (equivalent to murine L604) is associated with a higher risk for schizoaffective disorder (30). Further stabilisation of the A-B interface is mediated by conserved salt bridges spanning between the two CC3s (see R573 and R574 in protomer B binding to E619 and D579 in protomer A respectively, Supp Fig. 6a). The two independent interfaces between each A and B protomer contribute to an overall buried surface area of ∼4126Å^2^ (Fig. 2c), representing by far the biggest contributor to the stability of the tetramer folding.

#### DISC1 tetramer formation is underpinned by hydrophobic and charged interactions

Compared to the ‘monomer-monomer’ interface discussed above, the A-B dimers further combine together via the A-A interface by burying a relatively smaller surface area (∼1043Å^2^, Fig. 3a). Here, the two CC3s intersect at the point of the symmetry axis with a 30° angle (Supp Fig. 6b) which allows for an almost parallel, side to side, engagement of the two long α8 helices (Fig. 3b). Close inspection reveals an interesting partitioning of the interacting residues: hydrophobic interactions are found at the central part of the A-A interface, where the helices are closest to each other, involving the conserved residues L567, L571, L618 and F621 (Fig. 3b, right inset). Flanking those contacts are well-positioned electrostatic patches that, although reaching over a larger gap between the helices, still contribute through the bulk of ionic interactions to ‘dimer-dimer’ association (Fig. 3b, left inset).

**Fig 3.**
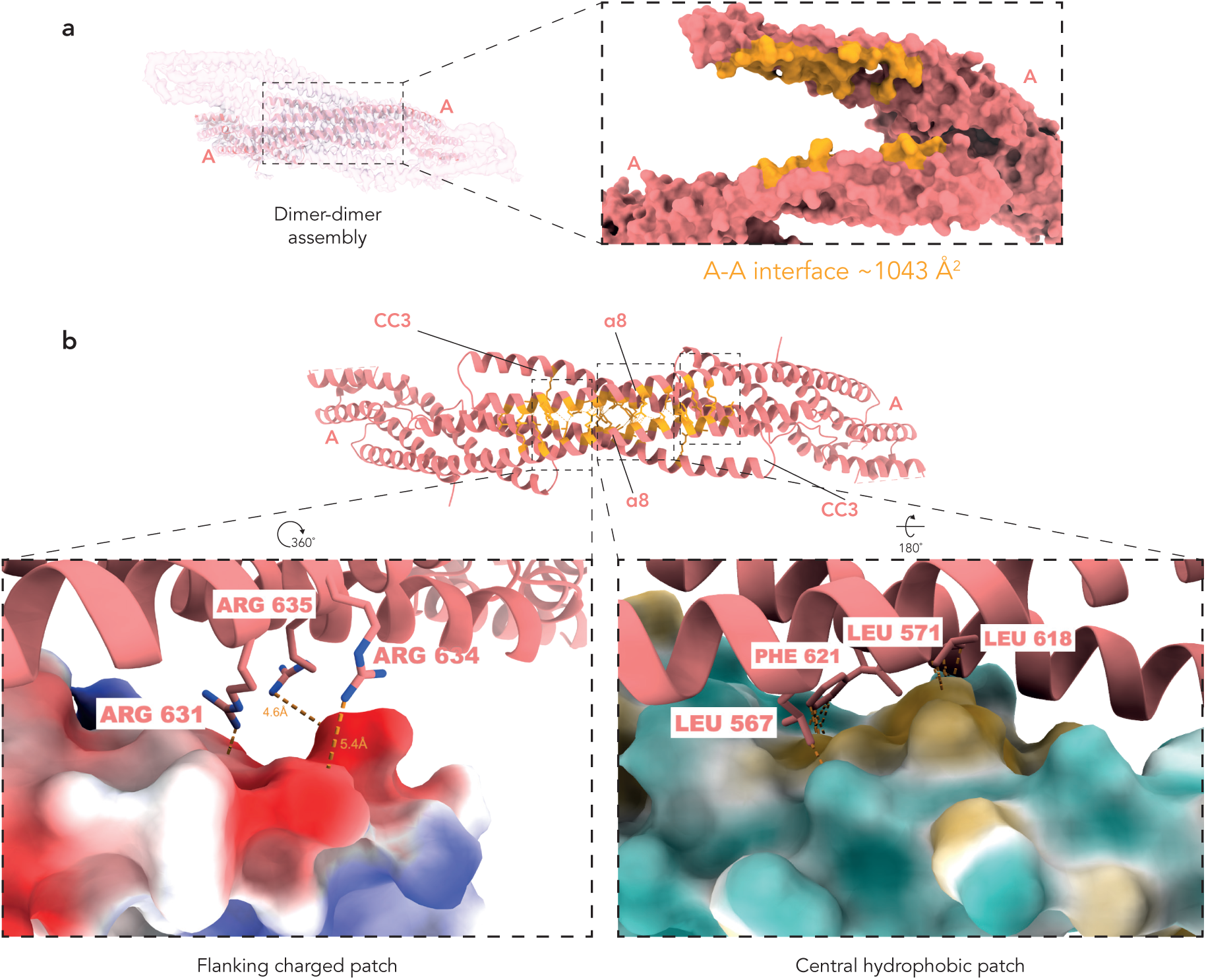
Both hydrophobic and ionic interactions contribute to tetramer formation. **a**: The two protomers A engaged at the dimer-dimer binding site are fitted into the EM map here. Like in Fig. 2c, left inset presents an exploded view highlighting the interface area in orange. Again, each monomer is coloured as in Fig. 1f. **b**: Overview of all contacting residues (in orange) at the A-A (‘dimer-dimer’) interface with the two CC3s that are involved indicated. The structure clearly shows an enrichment of contacts on the two opposing α8 helices. Below, left and right insets focus on the distinctive spatial organisation of interacting residues into a central hydrophobic core (right inset), which is then flanked by two electrostatic patches (left inset, only one of the two sites shown here for depiction clarity). The orientation of each inset view with respect to the reference structure above is also indicated.

It is likely that that further ‘dimer-dimer’ contacts occur along the sides of the cross-shaped DISC1 tetramer via the predicted CC2 coiled coils, for which a model could not be confidently built (Fig. 1c, Supp Fig. 6c). Indeed, the poorly resolved densities in these peripheral areas of the map suggest that CC2 is likely flexibly connected to the rest of the complex which is more rigid.

Overall, it is evident from our model that CC1 and CC3 are most instrumental to DISC1_core_ folding. Indeed, among the bundles of helices located away from these two domains (e.g. α3 packing against α6 and α1 against α9 respectively, Fig. 4a, right inset) none contribute to any of the protomer interfaces. The distribution of many buried hydrophobic residues within these helical bundles also do not seem to follow the heptad pattern found in genuine coiled coils. While both CC1 and CC3 do display the typical heptad repeats, they differ significantly in their length and arguably their structural bearing in DISC1 (Fig. 4a). CC1 only carries two heptads on each helix (Fig. 4b left) and is therefore most similar in size to a *bona fide* UVR domain. In comparison, CC3 is composed of many more heptads (Fig. 4b right) that extend well beyond the boundaries of a typical UVR motif. Given CC3 contributions to both A-A and A-B interfaces as well as the extent of hydrophobic residues buried in such a long coiled coil, this domain is arguably one of the most important structural elements contributing to a stable DISC1 fold. Testament to this notion, the hairpin loop region of CC3_UVR_ harbours the site (G598 in human DISC1) where protein truncation occurs as a result of the t(1;11) disease mutation (2,3). Our structure clearly suggests that the entire 80Å long CC3 would be completed abrogated by the truncation event (Fig. 4c). To experimentally validate the presumed impact of this disease mutation on DISC1 oligomerisation, we purified a murine DISC1_core_ mutant (DISC1 ΔN_mut B_) that is truncated after residue G595 (equivalent to G598 in human DISC1). Notably, this particular mutant form could only be isolated in solution when a N-terminal MBP tag was present. In agreement with our structural model, mass photometry analysis clearly indicates the absence of any tetramer assembly in DISC1 ΔN_mut B_ (Supp Fig. 5, upper plot, Table S4). Our result is very much similar to previous biochemical studies of the human t(1;11) mutant expressed recombinantly (31).

**Fig 4.**
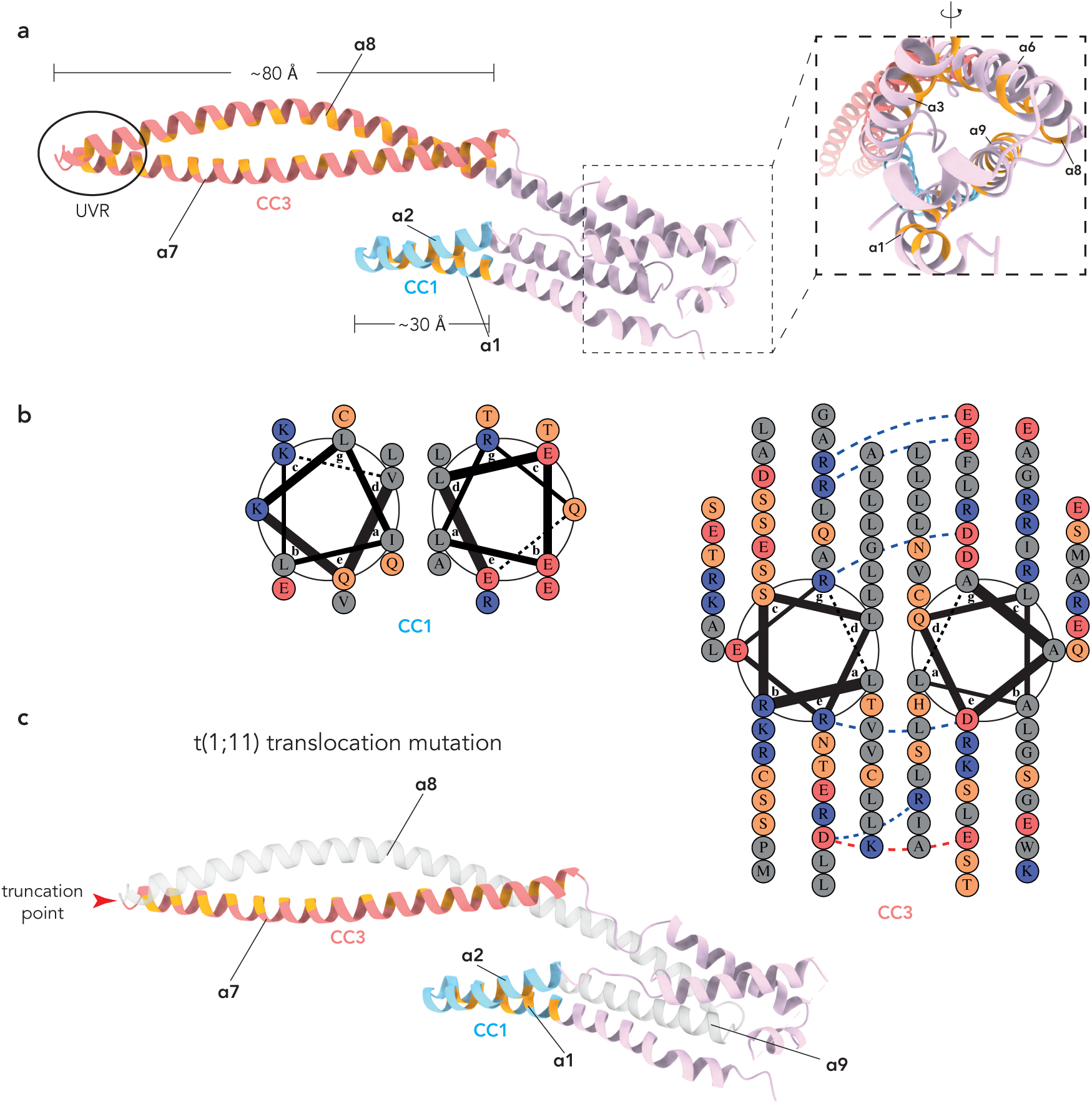
CC1 and CC3 are key structural elements of the DISC1 fold. **a**: Both CC1 and CC3 domains are color-coded as in Fig. 1a with their physical dimensions also annotated here. In particular, the hydrophobic residues occupying either the *a* or *d* position in the heptad repeats are shown in orange for both domains. In the case of CC3, the heptads can be seen to extend well beyond its UVR domain (black oval circle). The right inset shows that, beside CC1 and CC3, other helices in DISC1_core_ are also rich in hydrophobic residues, which facilitate their bundling together. The distribution of these residues does not seem to conform to a genuine coiled coil pattern. **b**: Helical wheel diagram showing the predominance of hydrophobic residues at positions *a* and *d* for both CC1 and CC3, thus confirming their status as *bona fide* coiled coils. The diagrams were made using the online tool DrawCoil 1.0. **c**: The diagram aims to illustrate the impact of the familial t(1;11) translocation mutation on DISC1_core_ folding. The red arrowhead pinpoints to the truncation site. The part of the protein that is lost as a consequence of the disease mutation is shown in semi-transparent light grey.

### DISC1 tetrameric core coordinates multiple partner proteins

#### A single DISC1 tetramer links multiple NDE1 partners

As evidenced by our structure, the numerous contacts at the two ‘monomer-monomer’ interfaces and the ‘dimer-dimer’ interacting site contribute to a combined buried surface area of ∼5169Å^2^ (not accounting for the dimerization pairs at CC2). Therefore, the extent of residue packing makes a strong argument for a constitutive tetrameric state of DISC1_core_ in solution. The high level of conservation observed for this region suggests that this fold has likely been preserved across evolution and may perform a functional role. We argue that a principal function of DISC1 is likely the coordination of multiple binding partners via tetramerization of its core. To test this hypothesis, we sought to reconstitute *in vitro* our DISC1 tetramer with its high affinity binder, NDE1 (9) and evaluate the complex stoichiometry. Importantly, we included in this assay a construct of DISC1 (denoted as DISC1 ΔN_WT_, residues 322-852) that contains both the core domain and the C-terminal NDE1 binding module, DISC1-C. Negative stain EM images of SEC-purified DISC1 ΔN_WT_-NDE1 complex (hereby abbreviated as DN_WT_, Supp Fig. 6d) clearly shows numerous instances where a single DISC1 tetramer is engaged with more than one molecule of NDE1 (Fig. 5a). Further quantitative analysis of this complex population by mass photometry (MP), revealed the presence of two distinct DN_WT_ species matching closely the molecular weight of a DISC1 tetramer carrying either one or two NDE1 homodimers (346 kDa and 429 kDa respectively, Fig. 5b).

**Fig 5.**
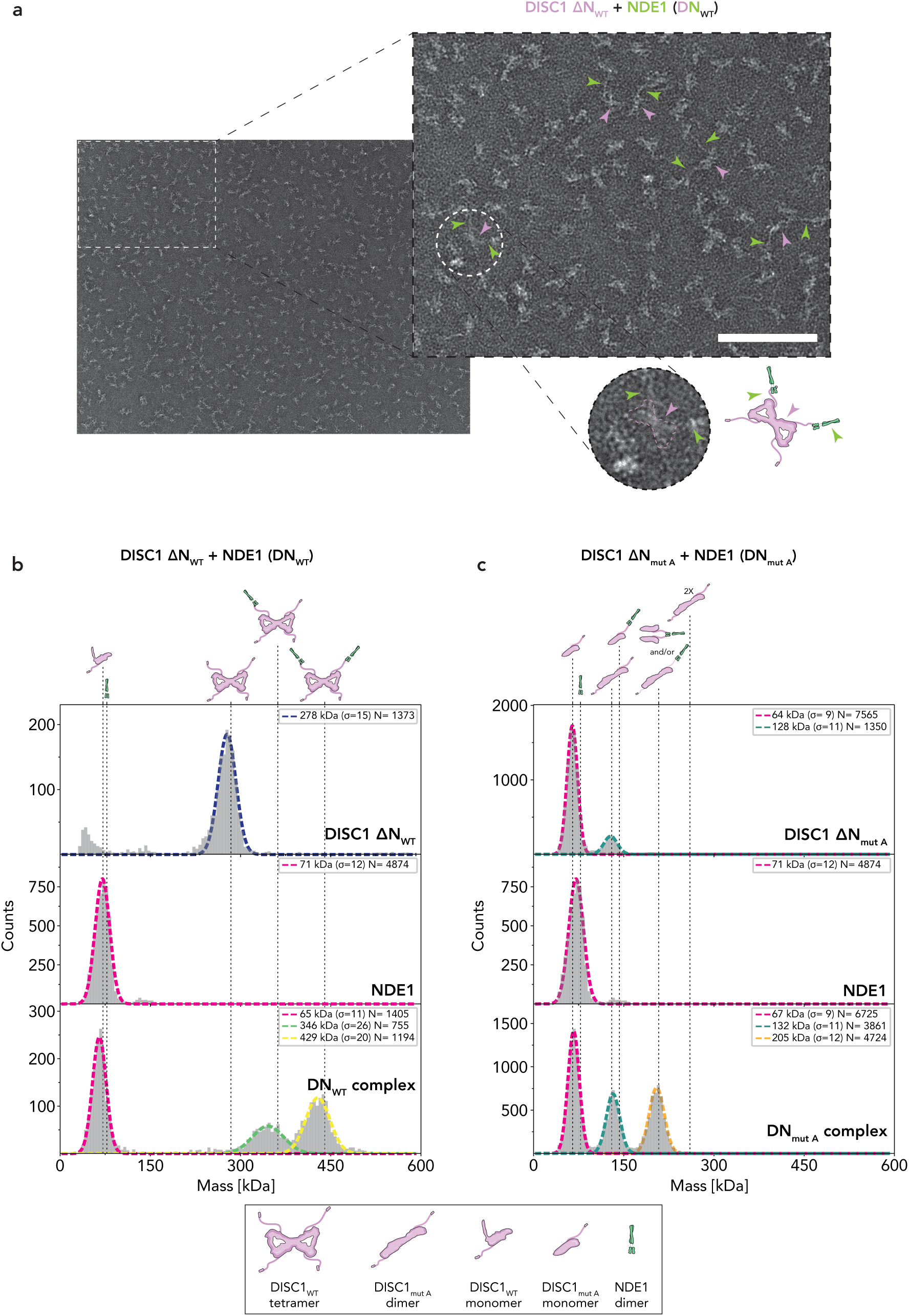
DISC1 tetramerisation enables coordination of multiple partner proteins. **a**: Negative stain EM micrograph showing *in vitr*o reconstituted and SEC-purified DISC1 ΔN_WT_-NDE1 (DN_WT_) complex. Right inset shows a close-up view of the protein particles on grids. Purple arrow heads point to DISC1 tetramers with their peculiar bow-tie like shape, while green arrow heads points to NDE1 molecules which are seen as filamentous-like particles due their well-known, extended coiled coil nature. The lower round inset provides an example of many instances where a single DISC1 tetramer coordinates more than one copy of NDE1. Here, the purple dashed contour highlights the boundaries of the DISC1 tetramer while the adjacent cartoon illustration depicts the overall DISC1-NDE1 arrangement observed. Scale bar = 50 nm. **b**: MP analysis of reconstituted DN_WT_ complex. Vertical dashed lines indicate the expected positions for various molecular species, each represented by a cartoon, according to their molecular weights (MW). Measured mean MW values for the Gaussian-fitted peaks (various colours) are given. The number of detected molecules contributing to each Gaussian distribution, and the corresponding standard deviation are also indicated. Top and middle plots show measurements of DISC1 ΔN_WT_ and NDE1 on their own respectively. While DISC1 ΔN_WT_ exists as a homotetramer (expected MW ∼284 kDa), NDE1 behaves as a homodimer (theoretical MW ∼78 kDa). Bottom plot shows the different species present in the reconstituted DN_WT_ sample. The 346 kDa and 429 kDa peaks clearly correspond to complexes where DISC1 is engaged with one or two copies of NDE1 homodimer respectively. **c**: As in b, MP measurements for DISC1 ΔN_mut A_ and NDE1 are shown in the upper and middle plots respectively. Note the appearance of a ∼205 kDa peak in the lower plot which likely correspond to a heterocomplex of DISC1 ΔN_mut A_-NDE1 (DN_mut A_). Either a dimer of DISC1 ΔN_mut A_ engaged with a single NDE1 dimer or two distinct DISC1 ΔN_mut A_ monomers coordinated by a NDE1 dimer or both those forms may be present here. The same type of cartoon illustration as in b is used.

#### DISC1 binding to NDE1 is independent of its tetramerization function

In accordance with DISC1 domain organisation, our cryo-EM reconstruction of DISC1 ΔN_WT_ revealed an almost indistinguishable resemblance to the DISC1_core_ tetramer (Supp Fig. 6e), confirming the notion that the addition of DISC1-C does not alter the core architecture. We wondered whether the inverse is also applicable, specifically if DISC1 tetramerization is dispensable for NDE1 binding. To this end, we abrogated DISC1 tetramer formation by both removing the CC2 domain (which as aforementioned could not be modelled confidently) and mutating key residues at the ‘dimer-dimer’ interface between the two protomers A in our structure (Supp Fig. 6f, Table S3). In support of our model, MP analysis confirms that this particular mutant (denoted here as DISC1 ΔN_mut A_) does not form any detectable tetramer but only exists as dimeric and monomeric species in solution (Fig. 5c, top plot, Table S4). Importantly, DISC1 association with NDE1 is not affected by the structural mutations, as evidenced by the emergence of a distinctive population, likely corresponding to a heterocomplex of DISC1 ΔN_mut A_-NDE1 (DN_mut A_, Fig. 5c, bottom plot) when these two proteins are reconstituted together *in vitro*.

## Discussion

Following decades of research interest in understanding the molecular architecture and function of the schizophrenia-associated protein DISC1, our study here reveals a surprisingly elaborate tetrameric fold of the largest, structured and evolutionarily well-conserved part of this protein. Previous pioneering work have highlighted the oligomerisation properties of various DISC1 regions and provided compelling evidence of dimeric and tetrameric states for individual structural elements such as CC1 and CC3 (32,33). Building on those earlier findings, our current structural data reveals a more comprehensive view of the interrelationship between various DISC1 domains in achieving a complex assembly. Thus, the structure presented here is most relevant in the context of the largest known DISC1 isoform (L, Uniprot Q9NRI5-1) which encompasses all of the domains discussed in this study. Nonetheless, it would be of great interest for future structural investigations to explore the potentially varied architectural landscape among other known DISC1 variants (34) and understand the possible roles played by previously identified oligomeric interactions in such context.

The idea of DISC1 serving as a pleiotropic molecular scaffold protein have been extensively reviewed before (1,15,35–37). The data presented in the current study provides an important structural basis for understanding DISC1 function as a multivalent hub. Indeed, we show that tetramerization enables DISC1 coordination of multiple copies of its client protein NDE1. Importantly, we also demonstrate that the capacity for DISC1 to form such a multivalent molecular platform is independent of its ability to interact with the partner. In support of this notion, we did not observe any structural changes in DISC1_core_ folding when the DISC1-C domain was present (Supp. Fig. 6e). The latter is also not exclusively targeted by NDE1 since other partners such as ATF4, a transcription factor, is known to share the same binding site (38). Indeed, the rather common coiled coil nature of these interactions suggest that DISC1-C may be a versatile binding module for other yet uncharacterised coiled coil-containing partner molecules. In addition to DISC1-C, the largely disordered DISC1-N has also been shown to harbour binding motifs for known binders (39,40). Therefore, it can be argued that, while DISC1_core_ may present a conserved, multivalent architecture regardless of the cellular context, the nature of DISC1 contribution at any given time likely depends on its subcellular localisation and hence the accessibility of certain partner proteins.

Such distinction in function between multimerization and partner binding is useful when assessing the potential implications of disease mutations and risk variants on DISC1 function. We argue that the severity of any mutational effect may correlate with the level of disruption to DISC1 tetramer folding. Among the many genetic mutations that have been reported for DISC1, the familial t(1:11) event is undoubtedly predicted to cause the most dramatic change in DISC1 structure. Indeed, as illustrated before, truncation of the protein chain at the CC3_UVR_ hairpin loop site will result into the complete loss of α8 helix and the consequent disassembly of CC3 (Fig. 4c). Since this domain is a key structural element contributing to both dimer and tetramer formation (Fig. 2 & 3), its abrogation will almost certainly abolish DISC1 multivalent architecture (Fig. 6). This scenario is strongly supported by our mutagenesis and mass photometry analysis (Supp Fig. 5). Consequently, any process that relies on DISC1 ability to bring different molecular players together is likely to be affected here. Not surprisingly, the t(1:11) mutation displays one of the strongest genetic linkage to a wide spectrum of psychiatric phenotypes that are not only confined to schizophrenia (4). In addition to loss of partner association, the t(1;11) mutation may also compromise DISC1 solubility by way of exposing normally buried hydrophobic residues that are prone to non-specific aggregation. Such notion is supported by previous reports of SDS-resistant species for the recombinantly expressed human t(1;11) mutant (31). Similarly, our DISC1 ΔN_mut B_ construct (murine DISC1_core_ harbouring the equivalent of the human t(1;11) truncation) could not be purified in solution without the addition of a bulky solubility tag like MBP. Interestingly, a previously reported frameshift mutation, targeting the very C-terminal end of DISC1 (5), has also been shown to lead to aberrant aggregations *in vitro* (32). In this case, the deletion caused by the frameshift event is also thought to induce disassembly of a coiled coil domain, namely DISC1-C. However, in contrast to t(1;11), this particular truncation is expected to only compromise binding to a select number of partners such as NDE1 and ATF4 without influencing DISC1 ability to tetramerise.

**Fig 6.**
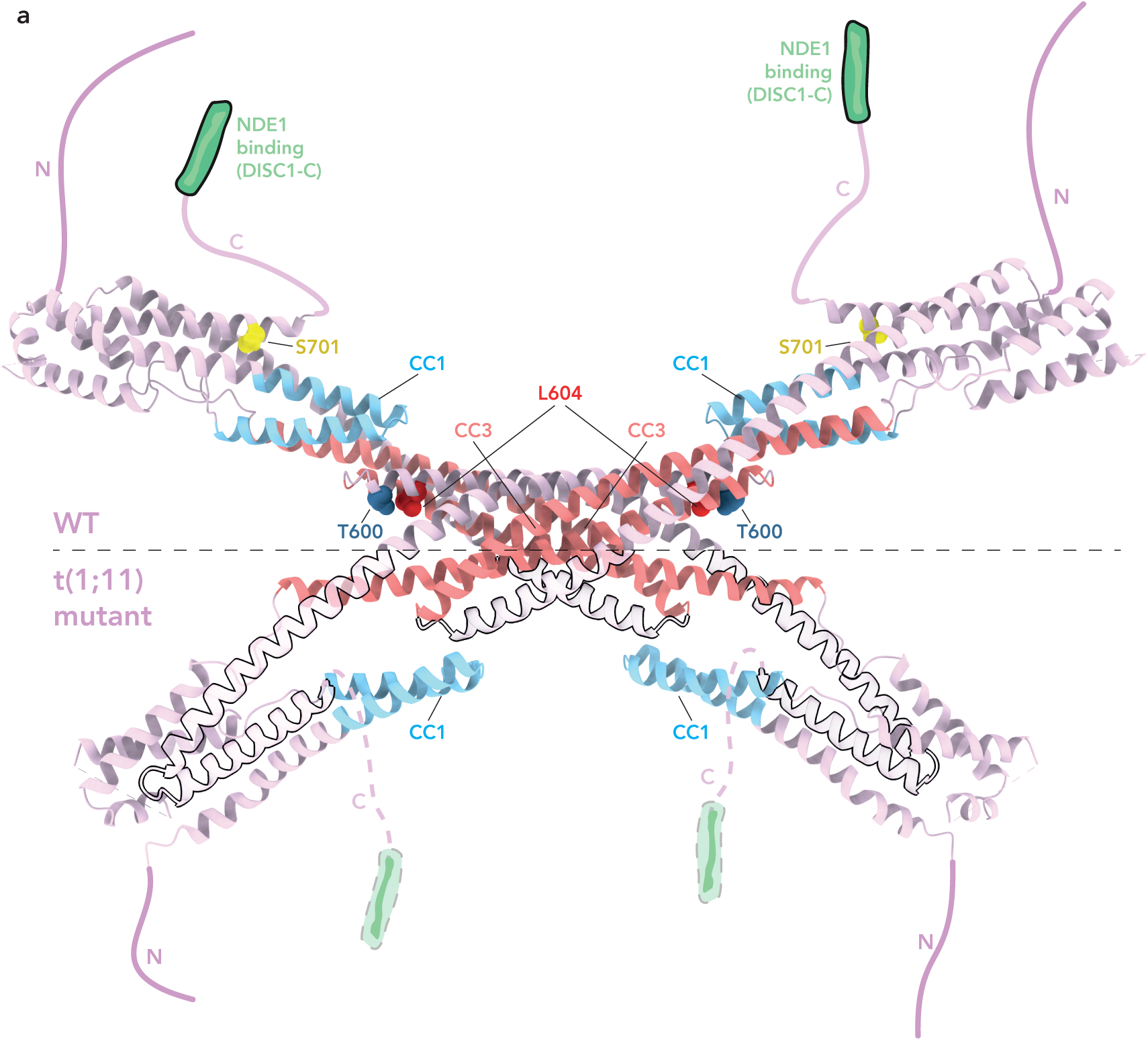
Effect of the t(1;11) disease mutation on DISC1_core_ structure. A side-by-side comparison between wild type (WT) DISC1 and the t(1;11) disease mutant in the context of the tetramer arrangement as well as its binding to NDE1 is shown here. Top half diagram illustrates flexible tethering of the C-terminal, NDE1-binding domain (DISC1-C, green cartoon representation) to DISC1_core_. The main structural elements CC1 and CC3 are highlighted here with the same colouring as in Fig. 1. Residues affected in three known disease risk variants, namely T600, L604 and S701 are also highlighted as sphere representations in dark blue, red and yellow respectively. These variants are likely to affect in subtle ways the overall stability of the tetramer folding while having a lesser impact on NDE1 association. For example, biochemical evidence indicate a propensity for the human S704C mutant to form higher-order oligomers while still maintaining NDEL1 binding (28). In contrast, as portrayed in the lower half of the diagram, the t(1;11) truncation affects both tetramerization (Supp Fig. 5, upper plot) and NDE1 interaction. Loss of the NDE1 binding module is represented here by faded dashed contour and dashed C-terminal linker. The region of the structure abrogated by the t(1;11) mutation is highlighted with a black contour and semi-transparent colouring. Note the deletion includes part of the CC3 as well as the above-mentioned SNP sites. Instead, both CC1 and CC2 (not shown here) remain intact.

Unlike the aforementioned deletions, the more common disease risk variants such as L607F and S704C are expected to generate subtler effects on DISC1 function. Mapping of those variants in the context of our tetramer model suggests a differential effect on the structure depending on their relative positions (Fig. 6). Indeed, both T600 (equivalent site of the human T603I SNP (7)) and L604 are within the CC3_UVR_ domain and therefore contribute to DISC1 oligomerisation. It is possible that the higher susceptibility to schizoaffective disorders demonstrated for those SNP variants are caused by increased instability of the tetrameric DISC1 structure. For example, the presence of a bulky aromatic side chain in L607F may result in steric clashes at the densely packed CC3_UVR_-CC1_UVR_ interface thus affecting the correct folding of CC3_UVR_ and/or dimerization. Future structural investigations should be able to clarify whether variants such as L607F induce changes in the molecular architecture of DISC1 and consequently affect proper coordination of binding partners. In contrast to T600 and L604, S701 is located at the periphery of the tetramer assembly and does not seem to participate directly to complex formation. Nonetheless, previous reports have highlighted a higher oligomerisation propensity for human S704C variant (28,33). It may be that those effects are underpinned by molecular interactions occurring through loop 2 which contacts with S704.

The extensive linker bridging DISC1_core_ and DISC1-C has attracted much research interest. It harbours the so called Lv splice site (34), which gives rise to an isoform with reduced affinity for NDE1 (41). Intriguingly, this splicing site also partially overlaps with known fibrillogenic sequences (42). It would be fascinating for future studies to decipher the interrelationship between splicing, fibrillation and NDE1 coordination within the more comprehensive context now provided by our structural model.

It is important to note that the murine DISC1_core_ shown in our structure shares only 63% sequence identity with the equivalent human domain (∼aa. 321-725) (43). However, the proteins display 90% hydropathy conservation, suggesting the same type of structure and function exist across this DISC1 region (Fig. 1a). In accordance with this notion, the same “bow tie”-like arrangement characteristic of our mouse DISC1 structure could be observed for a human equivalent construct under EM (Supp Fig. 6g). However, we were unable to obtain a high resolution cryo-EM map in this case.

In terms of evolutionary origin, it is interesting that the two CC1 and CC3 UVR motifs are the only homology domains been previously ascribed to DISC1 (21). Our study here confirms their importance as key building elements in the DISC1 fold. To our knowledge, this is the first known structural example of this ancient bacteria-derived domain contributing to the function of a mammalian neuronal protein. The degree of evolutionary conservation for the UVR dimer architecture between DISC1 homologues and the bacterial archetypes, to the extent of preserving the precise hydrophobic residues that mediate domain contact (including an ultra-conserved phenylalanine sitting within the loop, F598 in our DISC1 structure, Fig. 2b), is truly astonishing. Yet, the *E. coli* UvrABC endonucleases (22), the chloroplast Clp chaperone (25) and the mammalian DISC1 are found in seemingly unrelated cellular processes across different kingdoms. Similar to the DISC1 architecture described in this study, the UVR domains in bacterial and chroloplast proteins are flexibly linked to other functional domains that serve to confer specificity to their biological role. Therefore, it appears that the UVR domain has evolved as a context independent self-association hub to which other functional motifs can be tethered to.

The recruitment of ancient bacterial protein fold for the benefit of neuronal protein function in metazoans is not without precedence. Indeed, a number of neuronal synaptic proteins as well as surface adhesion receptors that have been implicated in mental illnesses are thought to have ancient evolutionary origin (44,45). In the case of DISC1, potential homologues have been identified in unicellular organisms such as the choanoflagellate *M.brevicollis* (21). Future studies of DISC1 homologues and their partner proteins in these simpler model systems will provide useful insights into the role played by the human protein in health and disease.

## Methods

### Vectors and cloning

Sequence encoding mouse DISC1_core_ residues (aa. 322-722) was cloned into a pET32MG vector containing N-terminal 6xHis affinity and GB1 solubility tags. Mouse DISC1 ΔN_mut B_ (aa. 322-595) was cloned into a modified pET32M vector that includes N-terminal MBP solubility and 6xHis affinity tags. All other mouse DISC1 ΔN constructs (aa. 322-852, including DISC1 ΔN_WT_, DISC1 ΔN_mut A_ and DISC1 ΔN_mut C_) were cloned into a modified pET32MG vector carrying N-terminal 2xStrepII affinity and GB1 solubility tags. Full-length mouse NDE1 and human DISC1_core_ (aa. 322-728) were cloned into a pET32M vector carrying N-terminal TrxA solubility and 6xHis affinity tags. All tags are cleavable by HRV-3C protease.

### Protein expression and purification

All recombinant proteins used in this study were expressed in *E.coli* BL21 CodonPlus or Rosetta 2 strains. Transformed cells were grown to an OD_600_ of ∼0.5 at 37°C before induction with 1 mM IPTG. Cells were then incubated at 16°C for 16 hrs before harvesting. Cells were re-suspended in 10 mL lysis buffer per each L of cell culture and lysed via sonication.

For the purification of the aforementioned StrepII-tagged mouse DISC1 constructs, the lysis buffer consisted of 50 mM Tris pH 8.0, 150 mM NaCl, 10% glycerol, 0.1% IGEPAL, 0.5 mM EDTA, 1 mM DTT supplemented with cOmplete protease inhibitor cocktail (Roche). Cleared lysate was incubated with StrepTactin affinity resin (IBA Life Sciences) in a gravity column and washed with a minimum of 40 column volumes (CV) of wash buffer (100 mM Tris pH 8.0, 150 mM NaCl, 1 mM EDTA, 1 mM DTT) and eluted in 1CV fractions with elution buffer (100 mM Tris pH 8.0, 150 mM NaCl, 1 mM EDTA, 5 mM desthiobiotin, 1 mM DTT), until no significant protein was detected in the elution by Nanodrop (Thermo Fisher Scientific) absorbance measurement at 280 nm. For DISC1 ΔN_mut WT_ and DISC1 ΔN_mut A_, peak elutions were further purified on a Mono Q 5/50 GL column (Cytiva) through 0.15-1M NaCl gradient in 50 mM Tris pH 8.0, 1 mM DTT. Serving as a final polishing step, both DISC1 ΔN_mut A_ and DISC1 ΔN_mut C_ were further separated on a Superdex 200 HiLoad 16/600 PG column (Cytiva) equilibrated in SEC buffer (50 mM Tris pH 8.0, 300 mM NaCl, 1 mM DTT). Elution fractions were concentrated to ∼1 mg/ml using Amicon Ultra Centrifugal filters (Merck Millipore). Aliquots were then flash-frozen in liquid nitrogen and stored at –80°C.

For the purification of 6xHis-tagged DISC1_core_ constructs (both human and mouse) as well as NDE1, the lysis buffer consisted of 25 mM sodium phosphate pH 7.4, 500 mM NaCl, 0.1% IGEPAL, 5 mM imidazole, supplemented with EDTA free cOmplete protease inhibitor cocktail (Roche). Cleared lysate were incubated with HisPur Ni-NTA affinity resin (Thermo Fisher Scientific) for 1 hour at 4°C and then washed in a gravity column with a minimum of 40 CV wash buffer (25 mM sodium phosphate pH 7.5, 500 mM NaCl, 20 mM imidazole). Protein sample were eluted in 1CV fractions with elution buffer (25 mM sodium phosphate pH 7.5, 500 mM NaCl, 250 mM imidazole), until no significant protein was detected in the elution by Bradford assay. Peak elutions were pooled and incubated for 16 hrs either in the absence (for DISC1_core_) or in the presence (for NDE1) of ∼75 μg of HRV-3C protease (purified in-house) per mL of elution, while dialysing against dialysis buffer (50 mM Tris pH 8.0, 150 mM NaCl, 1 mM DTT). Uncut DISC1_core_ and 3C-cut NDE1 were further purified on a Mono Q 5/50 GL column (Cytiva) as described for the StrepII-tagged DISC1 constructs. DISC1 peak elution fractions were again purified in a Superose 6 10/300 Gl column equilibrated in a SEC buffer containing 50 mM Tris pH 8.0, 150 mM NaCl, 1 mM DTT. Instead, NDE1 peak elutions were further cleaned in a Superdex 200 HiLoad 16/600 PG column (Cytiva) equilibrated in SEC buffer (50 mM Tris pH 8.0, 300 mM NaCl, 1 mM DTT). Elution fractions were then processed and stored in the same way as detailed above.

### Size-exclusion chromatography coupled to multiangle light scattering

100 μL of 1 mg/ml DISC1_core_ protein, for which the GB1 and 6xHis tags have been removed via 3C protease, was loaded onto a Superose 6 10/300 GL column (Cytiva) equilibrated in a buffer containing 50 mM Tris pH 8.0, 150 mM NaCl, 1 mM DTT on an AKTA FPLC system. This system was coupled to a light scattering detector (miniDAWN, Wyatt Technology) and a refractive index detector (OptiLab, Wyatt Technology). Data analysis was carried out using ASTRA 6 (Wyatt Technology).

### Cryo-EM sample preparation

To ensure optimal sample quality for cryo-EM study, thawed DISC1 aliquots were subjected to an additional size exclusion chromatography step on a Superose 6 3.2/300 Increase column (Cytiva) equilibrated in SEC buffer supplemented with 5 mM DTT and 0.4% CHAPS. The eluted peak fraction (∼2.8 mg/mL; shown in Supp Fig. 1b) was immediately used for grid freezing. 3 μL protein sample was applied onto Quantifoil holey carbon R1.2/1.3 300 copper mesh grid (Quantifoil) that has been cleaned in a PELCO easiGlow glow discharger (Ted Pella) at 15 mA for 90 s. The grid was blotted for 6 s with blot force 0 using a Vitrobot Mark IV (Thermo Fisher Scientific) at 21°C, 100% humidity, before immediately plunge-frozen in liquid ethane.

### Cryo-EM data collection and processing

For the mouse DISC1_core_ cryo-EM dataset, a total of 6,039 movies from three different data collection sessions were recorded in a Titan Krios G3i TEM (Thermo Fisher Scientific) operating at 300 kV, at a nominal magnification of 81,000X, corresponding to a pixel size of 1.06Å/pixel at the specimen level. Movies were taken using a K3 direct electron detector equipped with Bioquantum energy filter (Gatan) and applying a slit width of 20 eV. A total dose of ∼40 e/ Å^2^ was applied for each movie. All data processing, except for map sharpening, was carried out in the cryoSPARC software package (46). All movies were motion corrected using the patch motion correction job. CTF estimation was performed using the patch CTF estimation job. A subset of all motion corrected and CTF estimated micrographs (∼10%) were used for blob picking, followed by 2D template generation via one round of 2D classification job. The best 2D templates were then used as references for template picking all micrographs. Particles were extracted using a box size of 448 pix. An initial stack of 1,439,840 particles was sorted via four consecutive rounds of 2D classification. This resulted into a ‘cleaned’ stack consisting of 386,121 particles. These served to generate five independent 3D templates via the Ab initio job, followed by their 3D classification using the heterogenous refinement job. Particles contributing to the two maps highlighted by the red dashed box in Supp Fig. 2a were then merged and used to generate a high-resolution raw map via the non-uniform (NU) refinement job (47). The two half maps derived from NU refinement, were inputted into DeepEMhancer (48) to create a final, sharpened map.

### Cryo-EM model refinement

Part of the AlphaFold (49) predicted model for the mouse DISC1 homologue (Q811T9-F1-v4) encompassing residues that correspond to DISC1_core_ (aa. 322-722) was docked into the sharpened cryo-EM map as a rigid body in UCSF ChimeraX (50). Initial flexible fitting was manually performed via ISOLDE (51) with the guide of both raw and sharpened maps. The model was then refined using Phenix real-space refinement (52). Local manual refinements were again made in ISOLDE before this was inputted into Phenix for a final round of real-space refinement. The model was validated with MolProbity (53). Molecular graphics and analyses were performed with UCSF ChimeraX.

### Negative stain EM

Negative stain analysis was performed using continuous carbon 300 mesh copper grids (EM Resolutions). Grids were cleaned in a PELCO easiGlow glow discharger (Ted Pella) at 15 mA for 45 s. 5 μL drop of diluted protein sample (∼10 ng/μL) was applied onto the grid and incubated for 2 min. The sample was blotted away with filter paper (Whatman) and 5 μL of 2% uranyl acetate solution was immediately applied to the grid. This was allowed to incubate for 30 s before being completely blotted away with filter paper. The grid was then left to air dry for a minimum of 10 min before imaging under the microscope. Images were acquired using a Talos L120C (Thermo Fisher Scientific) operating at 120 keV. Images were recorded using a 4k x 4k Ceta camera at a nominal magnification of 36,000X corresponding to 4.03 Å/pixel at the specimen level.

For the human DISC1_core_ negative stain dataset, 50 micrographs were collected and processed in cryoSPARC (46). A total of 77,632 particles were selected using the template picker and sorted through two rounds of 2D classification without CTF correction.

### *In vitro* reconstitution of DISC1-NDE1 complex

Equimolar amounts of DISC1 (DISC1 ΔN_WT_ or DISC1 ΔN_mut A_) and NDE1 proteins were mixed together and dialysed against reconstitution buffer (25 mM HEPES pH 7.4, 0.1 M KCl, 1 mM DTT) at 4°C for 2 hrs. The reconstituted complex was resolved on a Superose 6 3.2/300 Increase column (Cytiva) equilibrated in the same buffer. Co-elution of DISC1 and NDE1 was confirmed by Coomassie-stained SDS-PAGE. The peak fraction was then used for subsequent mass photometry measurements.

### Mass photometry analysis

Mass photometry measurements and analysis were performed according to standardized protocol (54). Data were acquired by TwoMP instrument (Refeyn) using regular field of the view and acquisition time of 120 s. Contrast to mass ratio was estimated using protein Dynamin-1 ΔPRD, and integration time of 30 ms was used for movie processing. In the case of DN_WT_ and DN_mut A_ complexes, peak fractions eluted from the SEC column were measured directly within one hour. For the individual DISC1 or NDE1 samples, thawed aliquots were cleared by centrifugation before measurements. Fast manual dilution (between 20 to 30-fold) of the protein sample was performed by placing either 19 μL or 23.2 μL of buffer on a gasket and then pipetting 1 μL or 0.8 μL of protein respectively into the buffer droplet. Data acquisition was initiated immediately after mixing the well by pipetting up and down. For each DISC1 ΔN sample (WT and mutants), at least three independent measurements were performed. For each DN complex (WT and mut A), at least two independent measurements were performed.

## Data Availability

Atomic coordinates have been deposited in the PDB with accession numbers 9RUX. The cryo-EM density map has been deposited in the Electron Microscopy Data Bank under the accession number EMD-54277. Source Data are provided with this paper.

## Acknowledgments

We thank the staff at the Biological Cryo-EM Center, HKUST, in particular Dr. Yingyi Zhang who assisted with cryo-EM data collection. This facility is generously supported by a donation from the Lo Kwee Seong Foundation. We thank Dr. Joseph Caesar, and Dr. Edward Lowe at the COSMIC cryo-EM facility, University of Oxford, for support with data processing. We thank Dr. Clinton Lau and Dr. Giedrė Ratkevičiūtė, Biochemistry Department, University of Oxford, for access to reagents and assistance with protein expression. We thank Dr. Clinton Lau and Prof. Rob Klose, Biochemistry Department, University of Oxford, for insightful discussion of the study. We thank Dr. Miguel Berbeira Santana and Dr. Tomas Heger for critical review of the manuscript. J.C.Z. was supported by an IAS Junior Fellowship (HKUST Jockey Club). E.S. was supported by the Wellcome Trust (202827/Z/16/Z and 226647/Z/22/Z) and the EMBO Young Investigator Programme. J.K. was supported by UK Research and Innovation (UKRI) under the UK government’s Horizon Europe funding guarantee through project Marie Sklodowska-Curie Actions (MSCA) Postdoctoral Fellowship NanoMassCreator (101062868) EP/X025713/1. P.K. was supported by Engineering and Physical Sciences Research Council (EPSRC) Leadership Fellowship EP/T03419X/1. Y.F. was supported by a General Research Fund (16104518). M.Z. was supported by the RGC of Hong Kong and a Kerry Holdings Fellowship.

## Author Contributions

J.C.Z. designed and performed most of the experiments in this study including: protein expression and purification, cryo-EM data collection and processing, model building and biochemical assays. J.K. performed mass photometry data collection and analysis. Y.F. performed cloning and initial SEC-MALS experiment. K.e.O. assisted with model refinement. P.K. provided access to mass photometry equipment. M.Z. provided initial funding. E.S. provided funding for the study. J.C.Z. analysed all data and prepared the manuscript with the assistance of E.S.

## Materials and Correspondence

Data supporting the findings of this manuscript as well as constructs generated for this study are available from the corresponding author upon reasonable request.

Correspondence: Jin Chuan Zhou

