## Supplementary Information for "The schizophrenia associated protein DISC1 is a multivalent tetrameric hub of conserved ancient fold"

**Supplementary Figures**

a

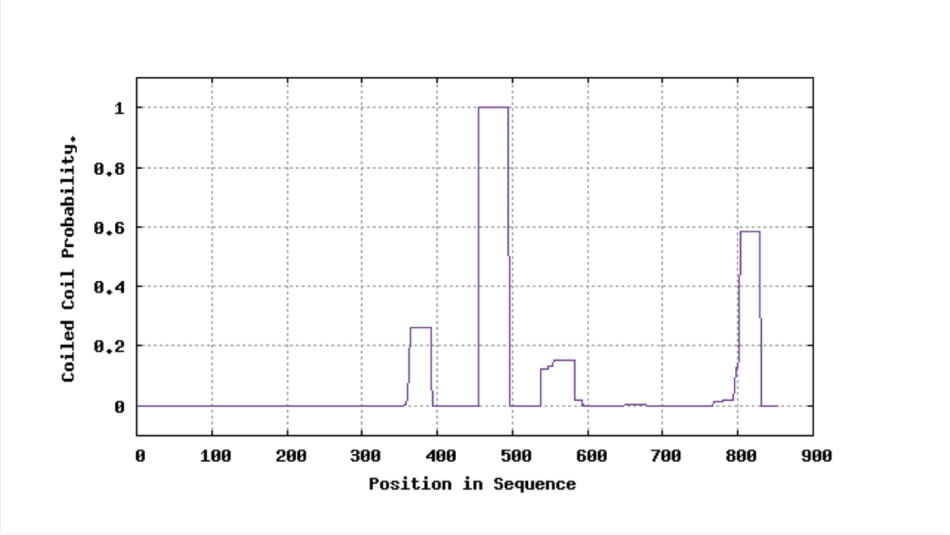


b

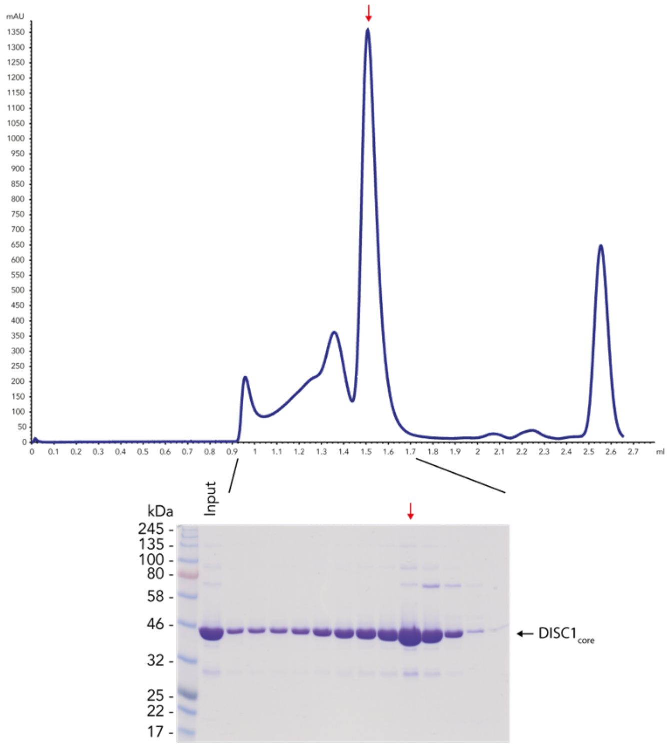


**Supp Fig 1. a**: MARCOIL prediction of coiled coil domains in the mouse DISC1 homologue. The first three peaks depicted in the diagram corresponds to CC1, CC2 and CC3 respectively within the DISC1 core region. **b**: SEC and SDS-PAGE analysis confirm the purification of a homogenous and monodisperse population of DISC1$_{core}$ sample. Red arrows indicate the elution peak in the chromatogram and the corresponding lane in the SDS-PAGE.

**a**

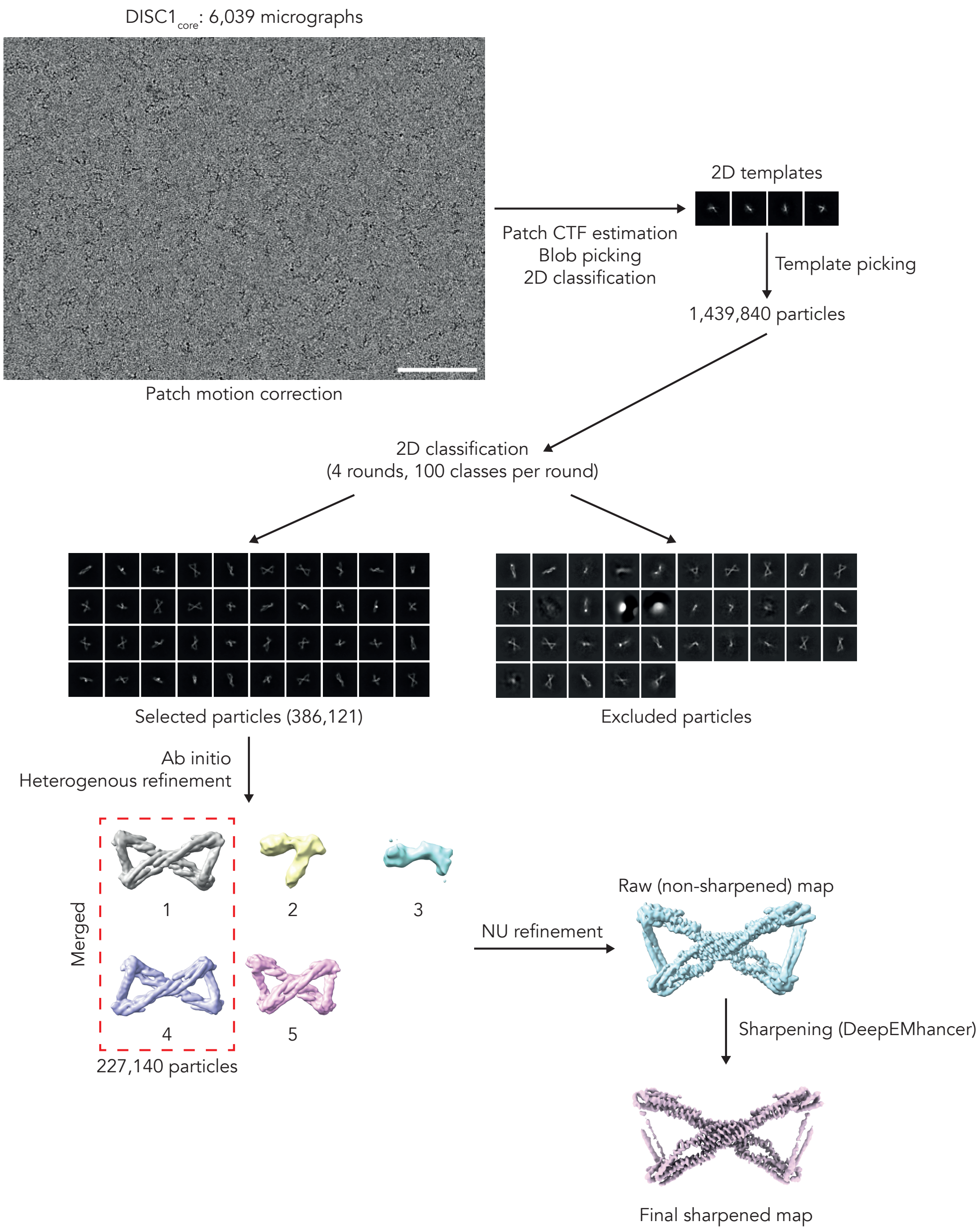

DISC1$_{core}$: 6,039 micrographs
Patch motion correction
Patch CTF estimation
Blob picking
2D classification
2D templates
Template picking
1,439,840 particles
2D classification
(4 rounds, 100 classes per round)
Selected particles (386,121)
Excluded particles
Ab initio
Heterogenous refinement
Merged
1
2
3
4
5
227,140 particles
NU refinement
Raw (non-sharpened) map
Sharpening (DeepEMhancer)
Final sharpened map

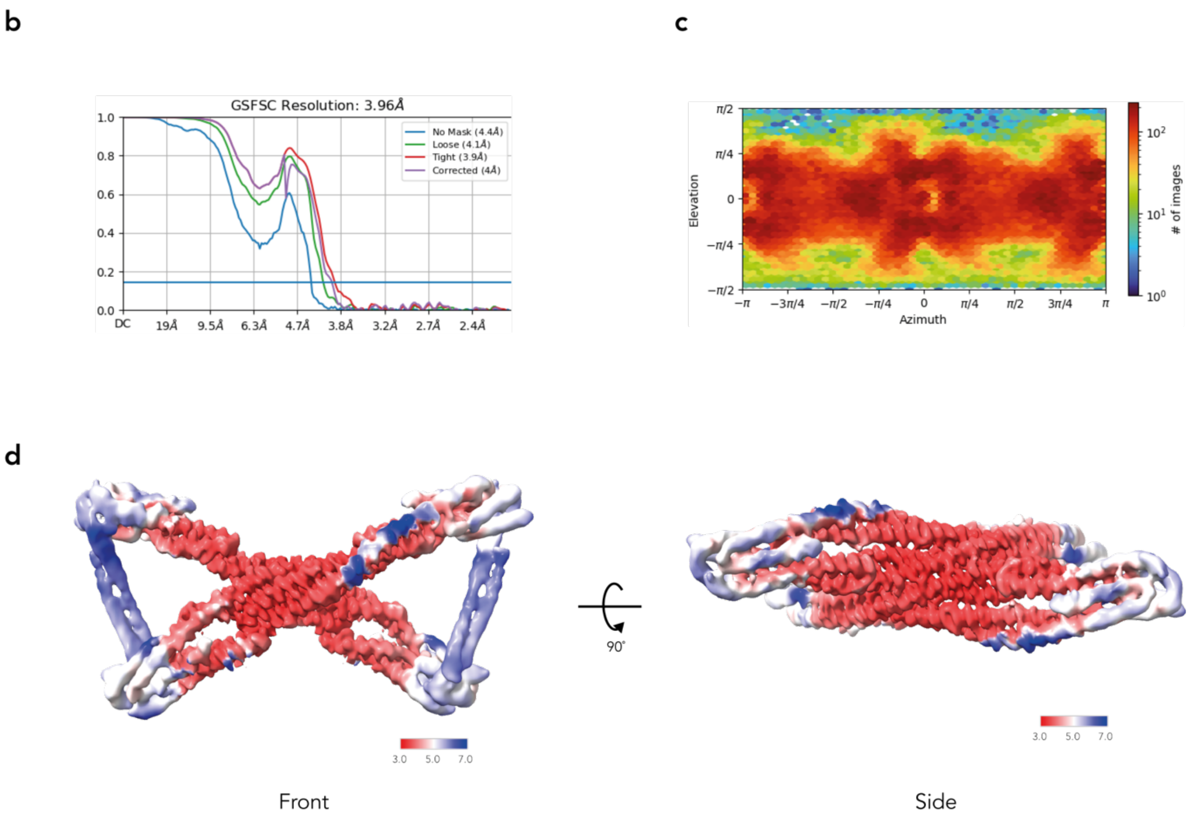


**Supp Fig 2. a**: Diagram illustrating the single particle cryo-EM data processing workflow. All data processing was carried out in the cryoSPARC software package except for final map sharpening, for which the DeepEMhancer program was used. At the top, a representative, non-dose weighted micrograph after the patch motion correction job is shown. Scale bar = 100 nm. Below, each data processing step, from 2D classification to final map sharpening, is detailed. Note 3D maps 1 and 4 resulting from the heterogenous refinement job represent the same DISC1 conformer. Therefore, particles contributing to those two maps have been pooled together prior to the NU refinement job. **b**: Gold standard FSC plot indicating the global resolution (cutoff=0.143) for the non-uniformly refined map in a. **c**: Euler angle distribution of all the particles that contributed to the refined map. **d**: Local resolution analysis of the refined map highlighting the higher resolution around the central region of $DISC1_{core}$ tetramer. Both front and side views are shown. Numbers displayed in colour key have units in Å.

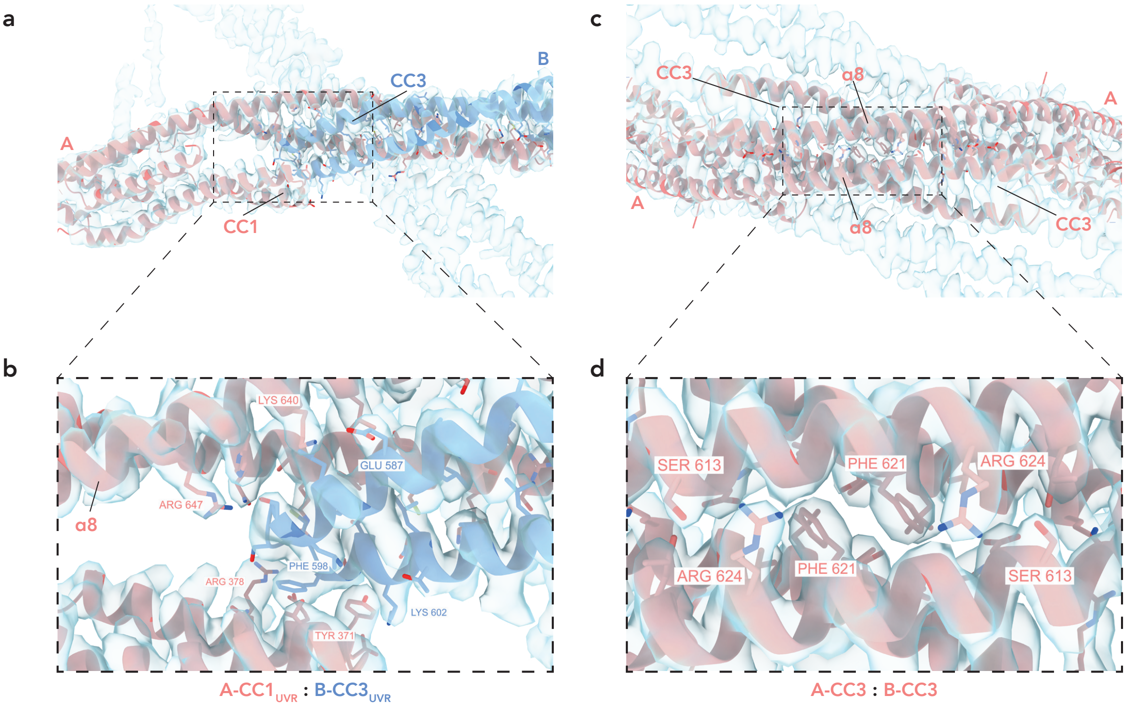


**Supp Fig 3. a**: Overview of the model-to-map fit at the A-B (‘monomer-monomer’) interface where the map is shown in semi-transparent light blue and the fitted A and B subunits are coloured as in Fig. 2. **b**: Close-up view of the correlation between model and map for the A-CC1$_{UVR}$ and B-CC3$_{UVR}$ interacting domains. For clarity, the fitting of only some interacting side chains are illustrated here. **c**: Overview of the model to map fit at the A-A (‘dimer-dimer’) interface with the same type of illustration as in a. **d**: Similar to b, a close up view of the fitting for some key interacting residues is shown here.

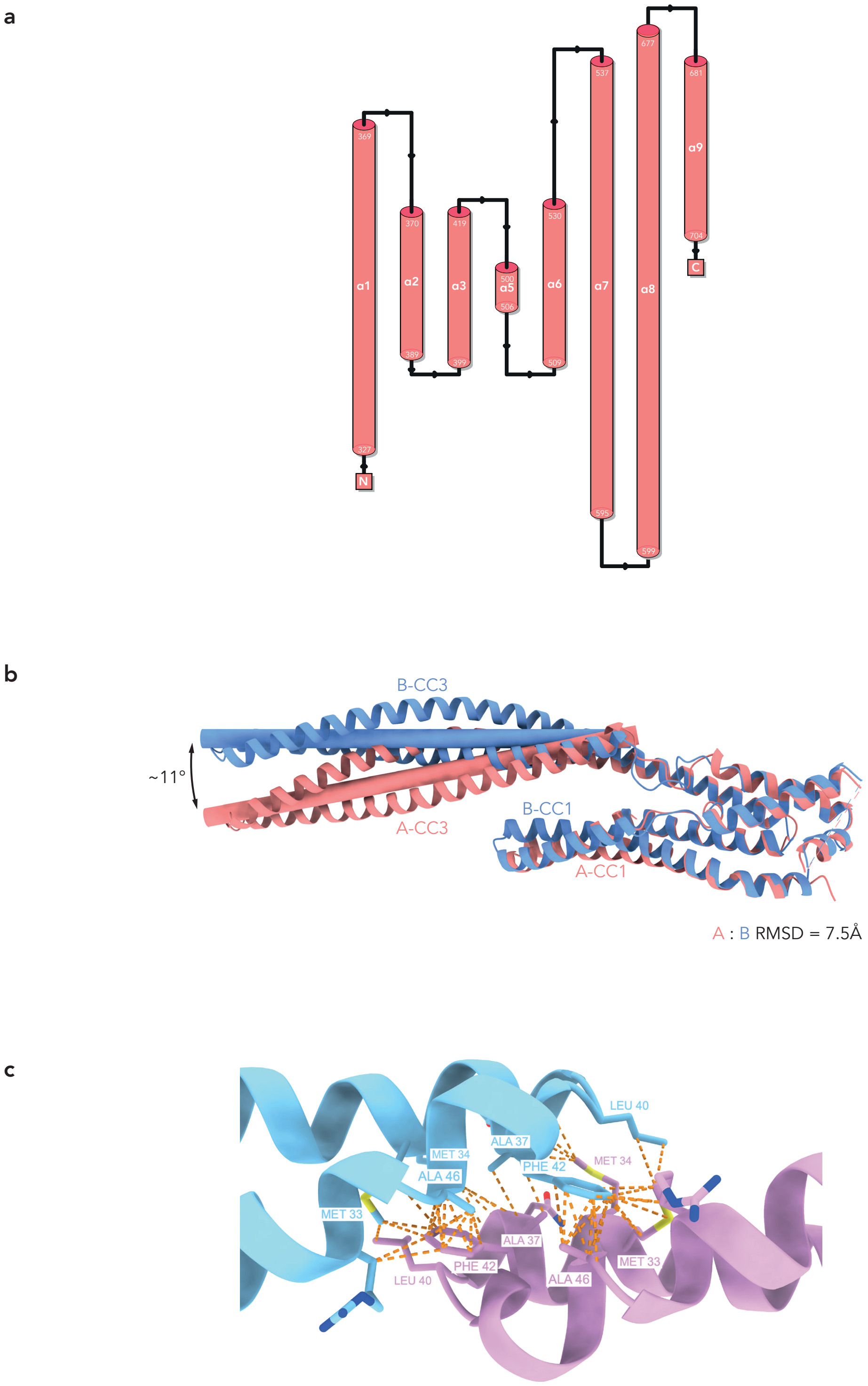


**Supp Fig 4. a**: Diagram delineating the topology of the eight helices that have been modelled based on our EM map. Only helix α4, which dimerises to form CC2, is not accounted for here since the CC2 densities in our map are too weak to be modelled with confidence. **b**: Structural superposition of A and B protomers reveal their asymmetric nature. The shift in CC3 tilt as well as the overall RMSD between the two monomers are indicated here. Slight changes in CC1 position can also be observed.

Each monomer is coloured as in Fig. 1f. **c**: UVR dimer of *E.coli* UvrB (pdb: 1e52) illustrating the conserved hydrophobic core at the dimerization interface.

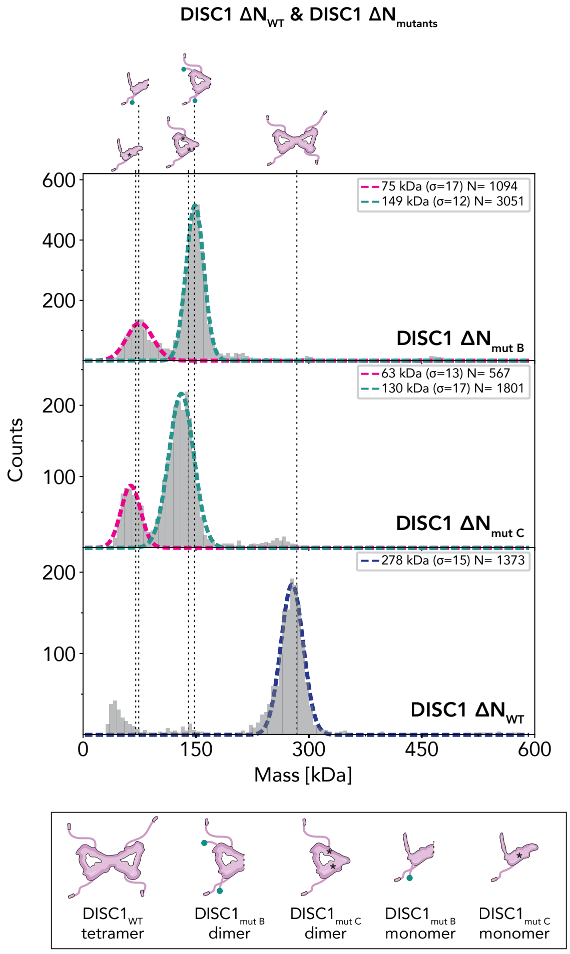


**Supp Fig 5.** Oligomerisation states of DISC1 $\Delta N_{mut\ B}$ and DISC1 $\Delta N_{mut\ C}$, as measured by mass photometry. Upper plot: DISC1 $\Delta N_{mut\ B}$ carries aa. 322-595, thus mimicking the truncation suffered by the human $DISC1_{core}$ due to the t(1;11) disease mutation. As evidenced here, only monomeric and dimeric species of this mutant form could be detected. Note that the presence of a N-terminal MBP tag (depicted by the green dot in the cartoon) is necessary to preserve protein solubility for this construct. Therefore, our *in vitro* result may not faithfully describe the actual molecular state of this particular mutant protein in the native cellular environment. Middle plot: despite DISC1 $\Delta N_{mut\ C}$ differing from its wild-type equivalent by only four residues located at the CC1-CC3 interface, this mutant is incapable of tetramer assembly. The black asterisk in the cartoon reflects the presence of point mutations. Lower plot: the same DISC1 $\Delta N_{WT}$ plot in Fig. 5b is shown here as a reference against the mutants shown above. Like Fig. 5, the number of detected molecules contributing to each Gaussian distribution, and the corresponding standard deviation are also indicated.

a

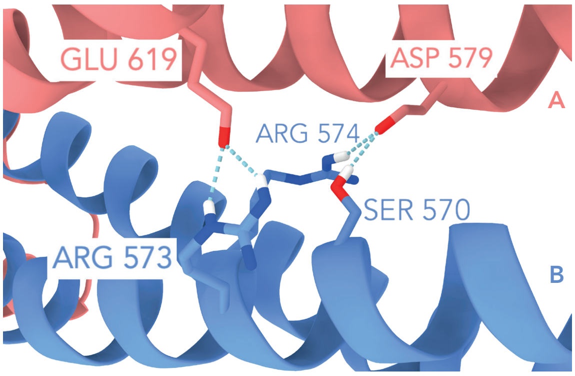
GLU 619
ASP 579
A
ARG 574
SER 570
ARG 573
B

b

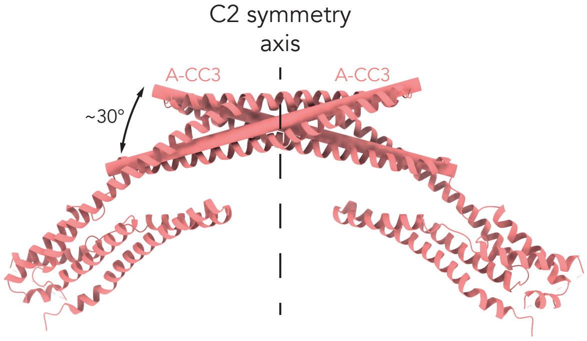
C2 symmetry axis
A-CC3
A-CC3
~30°

c

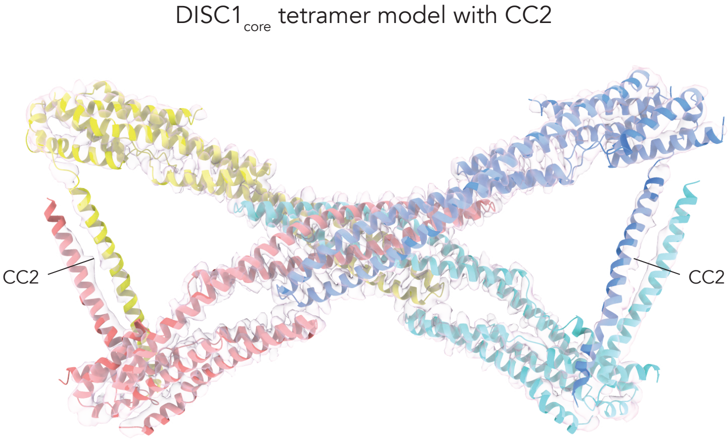
DISC1core tetramer model with CC2
CC2
CC2

d

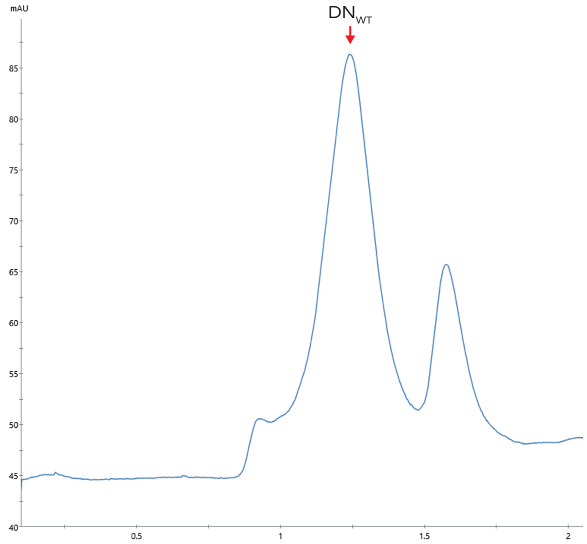


e

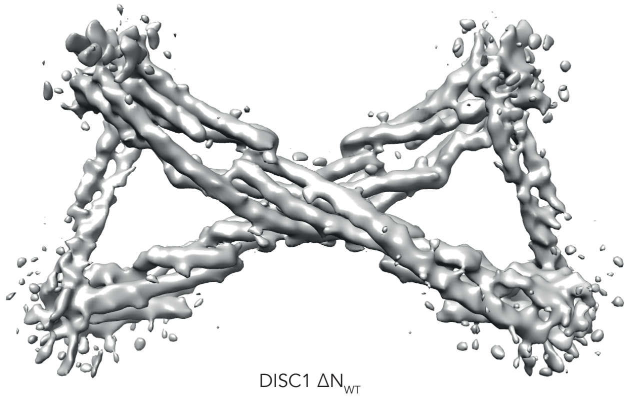


f

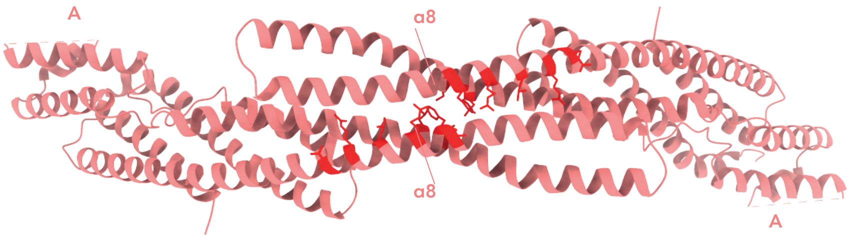


g

human DISC1 aa. 322-728

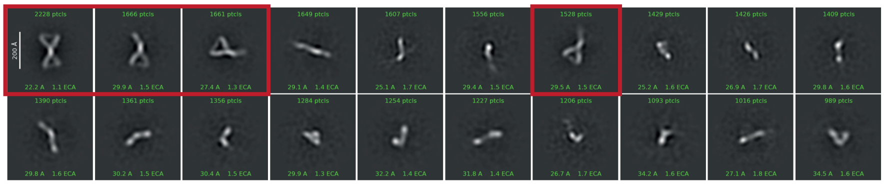

**Supp Fig 6. a**: Close-up view of two conserved salt bridges (R573-E619 and R574-D579) between the CC3s of opposing A and B monomers that contribute to the overall dimer stability. Each monomer is coloured as in Fig. 1f. **b**: Front view of the two protomers A bridging the 'dimer-dimer' interface. This clearly shows the two interacting CC3s crossing each other at about 30º angle. Each monomer is coloured as in Fig 1f. **c**: Atomic model of $DISC1_{core}$ where the two CC2 domains have been loosely fitted into the cryo-EM map. Note the exact positions of the four constituent α4 as well as their contacts could not be modelled with confidence. **d**: SEC chromatogram of reconstituted $DN_{WT}$ complex with the red arrow indicating the position of the $DN_{WT}$ elution peak. **e**: Cryo-EM reconstruction map of DISC1 $\Delta N_{WT}$ showing it adopts the same tetramer fold as $DISC1_{core}$. **f**: Overview of all the residues at the 'dimer-dimer' interface that have been mutated in DISC1 $\Delta N_{mut\,A}$. Here, we targeted residues that are part of the central hydrophobic cluster as well as those in the flanking charged patches. Each monomer is coloured as in Fig. 1f. Mutated residues are highlighted in red. **g**: Negative stain EM 2D classes of human $DISC1_{core}$ (aa. 322-728). Note the classes encircled in red display the same "bow tie"-like arrangement characteristic of mouse $DISC1_{core}$. As shown by the scale bar, these human DISC1 particles have a comparable size to the mouse DISC1 structure.

### Supplementary Tables

#### Table S1. Single-particle cryo-EM data collection and processing statistics

| | $DISC1_{core}$ |
|---|---|
| **EMDB ID** | |
| **Data collection and processing** | |
| Magnification | 81,000X |
| Voltage (kV) | 300 |
| Electron exposure ($e^-/Å^2$) | 40 |
| Defocus range (μm) | 1.25-1.75 |
| Pixel size (Å) | 1.06 |
| Symmetry imposed | C2 |
| Initial particle images | 1,439,840 |
| Final particle images | 227,140 |
| Map resolution | 3.9 |
| FSC threshold | 0.143 |

#### Table S2. Atomic model refinement statistics

| | $DISC1_{core}$ |
|---|---|
| **PDB ID** | |
| **Model refinement** | |
| Model resolution (Å) | 4.3 |
| FSC threshold | 0.5 |
| **Model composition** | |
| Chains | 4 |
| Non-hydrogen atoms | 9596 |
| Protein residues | 1216 |
| **RMSD deviations** | |
| Bond lengths (Å) | 0.005 |
| Bond angles (°) | 0.606 |
| **Validation** | |
| Clashscore | 4.69 |
| MolProbity score | 1.28 |
| Rotamers outliers (%) | 0.28 |
| **Ramachandran plot** | |
| Favoured (%) | 97.83 |
| Allowed (%) | 1.67 |
| Outliers (%) | 0.50 |

**Table S3. List of residues targeted in DISC1 ΔN$_{mut\ A}$ and DISC1 ΔN$_{mut\ C}$**

**DISC1 ΔN$_{mut\ A}$**

| aa. position | WT residue | Mutant residue |
|---|---|---|
| 451-494 | RRDWLIREKQRLQKEIEALQARMSALEAKEKRLSQELEEQEVLL | *deletion* |
| 567 | L | E |
| 570 | S | E |
| 620 | M | S |
| 621 | F | R |
| 624 | R | G |
| 628 | L | N |
| 631 | R | L |
| 634 | R | E |

**DISC1 ΔN$_{mut\ C}$**

| aa. position | WT residue | Mutant residue |
|---|---|---|
| 366 | V | E |
| 371 | Y | G |
| 593 | L | A |
| 598 | F | G |

**Table S4. Mass photometry statistics of DISC1 $\Delta N_{WT}$, $\Delta N_{mut\ A}$, $\Delta N_{mut\ B}$ and $\Delta N_{mut\ C}$**

**DISC1 $\Delta N_{WT}$**

| Tetramer MW (kDa) | Tetramer SD (σ) | Tetramer Counts (N) |
|---|---|---|
| 278 | 15 | 1373 |
| 271 | 15 | 1210 |
| 275 | 14 | 1261 |
| Average = 275 | | |

**DISC1 $\Delta N_{mut\ A}$**

| Monomer MW (kDa) | Monomer SD (σ) | Monomer Counts (N) | Dimer MW (kDa) | Dimer SD (σ) | Dimer Counts (N) |
|---|---|---|---|---|---|
| 64 | 9 | 7565 | 128 | 11 | 1350 |
| 65 | 10 | 8220 | 128 | 14 | 1571 |
| 62 | 12 | 6575 | 120 | 18 | 1245 |
| Average = 64 | | | Average = 125 | | |

**DISC1 $\Delta N_{mut\ B}$**

| Monomer MW (kDa) | Monomer SD (σ) | Monomer Counts (N) | Dimer MW (kDa) | Dimer SD (σ) | Dimer Counts (N) |
|---|---|---|---|---|---|
| 75 | 17 | 1094 | 149 | 12 | 3051 |
| 75 | 17 | 1229 | 145 | 12 | 3731 |
| 75 | 17 | 884 | 142 | 12 | 3175 |
| Average = 75 | | | Average = 145 | | |

**DISC1 $\Delta N_{mut\ C}$**

| Monomer MW (kDa) | Monomer SD (σ) | Monomer Counts (N) | Dimer MW (kDa) | Dimer SD (σ) | Dimer Counts (N) |
|---|---|---|---|---|---|
| 63 | 13 | 567 | 130 | 17 | 1801 |
| 68 | 17 | 990 | 130 | 16 | 3364 |
| 64 | 14 | 1048 | 132 | 15 | 3499 |
| Average = 65 | | | Average = 131 | | |

Statistics including the mean molecular weight (MW), standard deviation (SD) and the number of recorded counts in three repeat measurements for each DISC1 ΔN construct (WT and mutants) are given here. The average measured MW for each species is also indicated at the bottom. The expected MWs for the monomeric forms of DISC1 $\Delta N_{WT}$, $\Delta N_{mut\ A}$, $\Delta N_{mut\ B}$ and $\Delta N_{mut\ C}$ are 71 kDa, 65 kDa, 74 kDa and 70 kDa respectively.
